# Use of lignocellulosic residue from second-generation ethanol production to enhance methane production through co-digestion

**DOI:** 10.1101/2021.02.19.432018

**Authors:** Maria Paula. C. Volpi, Lívia B. Brenelli, Gustavo Mockaitis, Sarita C. Rabelo, Telma T. Franco, Bruna S. Moraes

## Abstract

This is a pioneer study evaluating the methane (CH_4_) production potential from residues of integrated 1^st^ (vinasse and filter cake) and 2^nd^ (deacetylation pretreatment liquor from straw) generation (1G2G) sugarcane biorefinery, providing a fully chemical characterization of them and their relation with the anaerobic digestion (AD) process. Small-scale assays provided fundamentals for basing the co-digestion optimization by assessing the optimal co-substrates synergistic conditions. Biochemical Methane Potential (BMP) tests showed co-digestion enhanced CH_4_ yield of isolated substrates, reaching up to 605 NmLCH_4_ gVS^-1^. The association of vinasse and deacetylation liquor as co-substrates increased the BMP by ~38% mostly by nutritionally benefiting the methanogenic activity. The kinetic analysis confirmed that the deacetylation liquor was the co-substrate responsible for improving the CH_4_ production in the co-digestion systems due to the highest CH_4_ conversion rate. The alkaline characteristic of the liquor (pH~12) also prevented alkalizing from being added to the co-digestion, an input that normally makes the process economically unfeasible to implement on an industrial scale due to the large quantities required for buffering the reactor. The filter cake had the lowest BMP (262 NmLCH_4_ gVS^-1^) and digestibility (≤ 40%), further limited by the required stirring to improve the mass transfer of biochemical reactions. The present study drives towards more sustainable use of vinasse, the most voluminous waste from the sugarcane industry, and lignin-rich residues derived from pre-treatment alkaline methods, aiming at an energy-efficient utilization, by at least 16% when compared to the traditional vinasse AD. The experimental and modeling elements from this work indicated the lignin-rich liquor is the main responsible for putting the co-digestion as a disruptive technological arrangement within the 1G2G sugarcane biorefineries, reinforcing the biogas production as the hub of the bioeconomy in the agroindustrial sector.

## 1. INTRODUCTION

Anaerobic digestion (AD) is an attractive process for managing liquid and solid organic waste that allows energy recovery through biogas, rich in methane (CH_4_). Organic matter conversion occurs by the activity of microbial consortia in a finely-tuned balanced ecosystem. Digested material i.e. digestate can also be exploited as a value-added by-product for agriculture [1, 2]. This biotechnological process is part of the current global context of searching for available residual substrates aligned to the diversification of product generation.

Despite all scientific growth in this area, gaining more knowledge based on innovative issues to comprehensibly investigate interactions between technological and fundamental bioprocess limitations entails optimizing CH_4_ generation. For example, the availability of biodegradable fractions in the substrates from the sugar-energy industry (related to AD with consequent CH_4_ production) still represents a bottleneck for this scientific field [3]. Insufficient knowledge on the principles and operation of AD bioreactors fed with such substrates often results in failed applications in Brazilian sugarcane mills. On the other hand, regarding pre-treatment processes for lignocellulosic biomass to obtain hexose and pentose fractions for other bioprocesses, as in the case of 2G ethanol production, enormous advances in fundamental and technological aspects can be found in the literature [4, 5].

Some by-products from the sugarcane agroindustry are already considered raw materials for the recovery and generation of value-added products [6]. Vinasse generated from ethanol distillation is commonly directed to sugarcane culture as liquid-fertile. For each liter of alcohol produced, approximately 10 L of vinasse are generated, and its composition is 0.28-0.52 g. L^-1^ of nitrogen (N), 0.11-0.25 g L^-1^ of phosphorus (P), 1.0-1.4 g L^-1^ of potassium (K), and 20-30 g L^-1^ of Chemical Oxygen Demand (COD) [7, 8]. Sugarcane bagasse, traditionally used in energy generation in Combined Heat and Power (CHP) systems, can be used as a substrate to produce 2G ethanol and other added value by-products [9]. Sugarcane straw, also considered a potential organic source, has become available as lignocellulosic biomass since the progressive introduction of mechanical harvest without burning procedures in Brazil [10]. In addition to being left in the field for agricultural reasons, straw can be used as feedstock for thermochemical or biochemical conversion processes, which makes it feasible to incorporate it into a biorefinery. Sugarcane straw has a similar chemical composition to bagasse in terms of the major components of biomass: cellulose (30-40% w/w), hemicelluloses (20-30% w/w), and lignin (15-30% w/w) [11]. This biomass can be converted into value-added products as biofuels, after pretreatment methods and multienzyme complexes to liberate sugars Among the diversity of methods that have been researching aiming at technological process improvements, Brenelli et al. [12] recently reported a promising alkaline pre-treatment of sugarcane straw by deacetylation, in which acetic acid is removed as it is an inhibitor for microorganisms in fermentation processes, and thus, xylo-oligosaccharide (XOS) are recovered for being fermented to ethanol. Filter cake, another organic solid byproduct, is generated from the filtration in rotary filters after cane juice clarification processes, presenting concentrations of 140-169 g kg^-1^ of lignin, 171-184.6 g kg^-1^ of cellulose and 153-170 g kg^-1^ of hemicellulose [3, 13]. It has been used in intrinsic steps at the plant (improvements in permeability during sucrose recovery in the rotary filter) [14] and as a source of nutrients for the soil [15]. Non-controlled digestion of such waste in the fields may lead to the release of large amounts of CH_4_, which may hinder the positive effect of bioenergy utilization on climate change mitigation [13].

The economic profitability of biorefineries can be supported by the integrated production of low-value biofuels [16]. In this context, co-digestion of residues can optimize CH_4_ production by providing and balancing macro and micronutrients for the AD process. It may also be the best option for substrates that are difficult to degrade. This appears to be the case for residues from ethanol production from the processing of lignocellulosic biomass, normally recognized as complex substrates for AD [7]. In addition to intrinsic improvements in the biological process (e.g. upgrading biogas production; better process stabilization by providing synergistic effects within the reactor; increased load of biodegradable organic compounds), the economic advantages of sharing equipment and costs are also successful [17]. Janke et al. [18] showed that co-digestion of filter cake with bagasse would produce 58% more biogas compared to large-scale filter cake mono-digestion. However, there are still gaps in the literature concerning the use of lignocellulosic residues from 2G ethanol production as cosubstrates.

The biodegradation capacity of residues can be assessed by Biochemical Methane Potential (BMP) assays. This approach shows the maximum experimental potential to convert the organic fraction of the substrates into CH_4_. Specific conditions in AD can also be evaluated: substrate sources (exclusive or blend proportions), temperature, nutrients, buffering, source of inoculum, among other factors. The BMP is the most used methodology by academic and technical practitioners to determine the maximum CH_4_ production of a certain substrate [19, 20].

This work aimed to determine the BMP of the main residues from 1G2G sugarcane biorefinery – vinasse, filter cake, and deacetylation liquor (waste stream from the pretreatment of sugarcane straw for the 2G ethanol production) – in assays as isolated substrates and as different blends of co-substrates for biogas production. The kinetic modeling from the experimental BMP data was further performed. The prior characterization of the substrates (in terms of nutrients) and its relation with the BMP results allowed to investigate the synergistic effects of the co-digestion on CH_4_ production, which was further proved by the kinetic analysis. The BMP and kinetics evaluation accounted for the CH_4_ production from the isolated substrates and from their co-digestion in different combinations.

## 2. METHODOLOGY

### 2.1 Substrates and inoculum

Vinasse and filter cake from a 1G sugarcane ethanol production process were obtained from Iracema Mill (São Martinho group), Iracemápolis municipality, São Paulo state, Brazil. The deacetylation liquor was obtained from a mild alkaline pretreatment of sugarcane straw under optimal conditions to remove acetate and lignin determined previously [12]. Briefly, 316L stainless steel batch reactor of 0.5 L capacity was filled with 20.0 g of raw sugarcane straw (dry basis) and NaOH aqueous solution at 8% w/w in 10% (w/w) of the final solid loading and incubated at 60°C for 30 min. Afterward, the reactor was immediately cooled in an ice bath and the liquid fraction separated from the solid fraction through a muslin cloth stored at 4°C until further use. The compositional analysis determined according to the NREL/TP-510-42623 procedure [21], showed the deacetylation liquor was mainly composed of acetate, soluble lignin, and lignin-derived compounds and extractives.

This study compared two different inocula to perform the anaerobic codigestion of sugarcane processing residues. In Experiment 1, the inoculum was obtained from the sludge of a mesophilic reactor (BIOPAQ^®^ICX - Paques) installed at Iracema Mill from São Martinho group (22°35’17.6”S 47°31’51.5”W) treating sugarcane vinasse. Experiment 2 used an anaerobic consortium from the sludge of a mesophilic Upflow Anaerobic Sludge Blanket (UASB) reactor treating poultry slaughterhouse wastewater from Ideal slaughterhouse, at Pereiras municipality, São Paulo state, Brazil (23°05’10.5”S 47°58’58.9”W).

All microorganisms used in this study are registered in the Brazillian Government’s system SISGEN (access number A5E04AF), accordingly with Brazillian law for accessing genetic resources and associated traditional knowledge.

### 2.2 Biochemical Methane Potential (BMP) of substrates

Theoretical Biochemical Methane Potential (TBMP) of filter cake was based on the Buswell equation (Equation 1) for solid substrates. TBMP of deacetylation liquor and vinasse was calculated from their organic matter concentration and VS content, as depicted by (Equation 2) for liquid substrates [19].

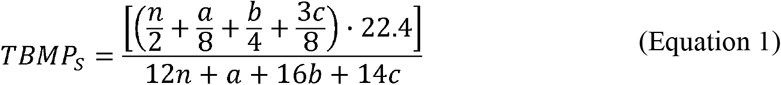

Where TBMP_S_ is the theoretical biochemical methane potential for solid substrate (NmLCH_4_ gVS^-1^), 22400 mL mol^-1^ is the molar gas volume in the standard temperature and pressure (STP, 273 K, and 1 bar). a, b, c and *n* are the molar content of the substrate for hydrogen, oxygen, nitrogen, and carbon, respectively.

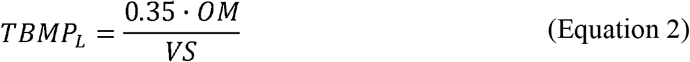

Where TBMP_L_ is the theoretical biochemical methane potential for liquid substrate (NmLCH_4_ gVS^-1^), 350 NmL is the theoretical CH_4_ yield of 1 g COD at STP [22], VS is the volatile solids of the substrate (g mL^-1^) and OM is the organic matter concentration, in terms of COD (gO_2_ mL^-1^).

BMP tests were performed to determine the biodegradability (BMP/TBMP) of crude substrates and their experimental potential for CH_4_ production following the protocol of Triolo et al. [23] and the VDI 4630 methodology [24]. All experiments were conducted in triplicates of single batches using 250 mL Duran^®^flasks as bioreactors closed with a pierceable isobutylene isoprene rubber septum, and stored in an Ethickthecnology (411-FPD) incubator at thermophilic condition (55°C) as vinasse leaves the distillation columns at 90°C and thus would have lower (or none) energy expenditure for cooling it to mesophilic conditions. As mesophilic sludges were used in thermophilic tests, the previous acclimatation of inocula was carried out for avoiding thermal shock to the microbial community. The temperature was gradually increased every 5 degrees until it reached 55°C, which was kept for a week before the beginning of the tests, as already demonstrated in the literature [25]. The experiment was 2:1 inoculum to substrate ratio (w/w, in terms of Volatile Solids-VS) added to each flask, thus ensuring excess of inoculum to consume all the organic matter of the substrate and achieving its maximum experimental CH_4_ production. The pH of solution flasks was corrected to neutrality by adding solutions of NaOH (0.5 M) or H_2_SO_4_ (1 M) when necessary. Nitrogen (N_2_) gas was fluxed into the liquid medium for 10 min and into the headspace for 5 min after closing the flasks. The headspace was kept at 40%. Biogas was collected from the headspace over the days by using a Gastight Hamilton Super Syringe (1L) through the flasks’ rubber septum. The measured biogas was corrected for a dry gas base by excluding the water vapor content in the wet biogas. The pressure and temperature for one-liter gas were corrected to normal (NL) following the standard temperature and pressure (STP) conditions (273 K, 1,013 hPa).

BMP was calculated through the average value of the replicates obtained at the end of each batch experiment, according to the traditional methodology [24] (Equation 3), and also through the kinetic modeling as the approach suggested by Filer et al. [26], as this latter considers the trending of the values and the fundamental parameters of the process. Both BMP results were compared and used to calculate the biodegradability.

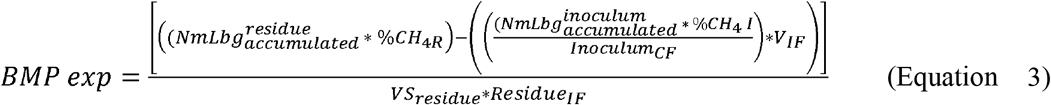

Where *BMP exp* is the biochemical methane potential of each residue (NmLCH_4_ gVS^-1^), 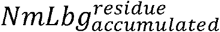 is the accumulated production of biogas from the residue (NmL), %CH_4 R_ is the CH_4_ content from biogas of the residue, 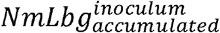 is the accumulated biogas production of inoculum (NmL), %CH_4 I_ is the CH_4_ content from biogas of the inoculum, *Inoculum_CF_* is the volume of inoculum in a flask of inoculum control (negative control) (mL), V_IF_ is the volume of inoculum added in each flask (mL), VS_residue_ is the number of volatile solids from the residue (gVS - g mL^-1^) and Residue_IF_ is the amount of residue added in the flask (mL or g).

Analysis of variance (ANOVA) was used to identify the existence of significant differences between the treatments and the Tukey test (p < 0.05) was performed to group BMP data. These analyzes were performed by Microsoft Excel version 12.

### 2.3 Experimental arrangement

Two rounds of BMP tests were performed. Experiment 1 assessed the inoculum from vinasse treatment (Section 2.1) and equal percentages (in VS terms) of substrates for the co-digestion test. Experiment 2 assessed the inoculum from poultry slaughterhouse waste treatment (Section 2.1) and the co-digestion conditions were expanded. The proportions of inoculum/substrate added in each flask were the same for both rounds of experiments (2:1 in terms of VS), as mentioned in section 2.2. Two types of control were used in both experiments: positive and negative. Microcrystalline cellulose (Sigma-Aldrich Avicel^®^PH-101) was used as a positive control (+). Negative control (-) was conducted only with each inoculum. Digestion was terminated when the daily production of biogas per batch was less than 1% of the accumulated gas production. The experimental design of both experiments is described in Table 1.

**Table 1.**
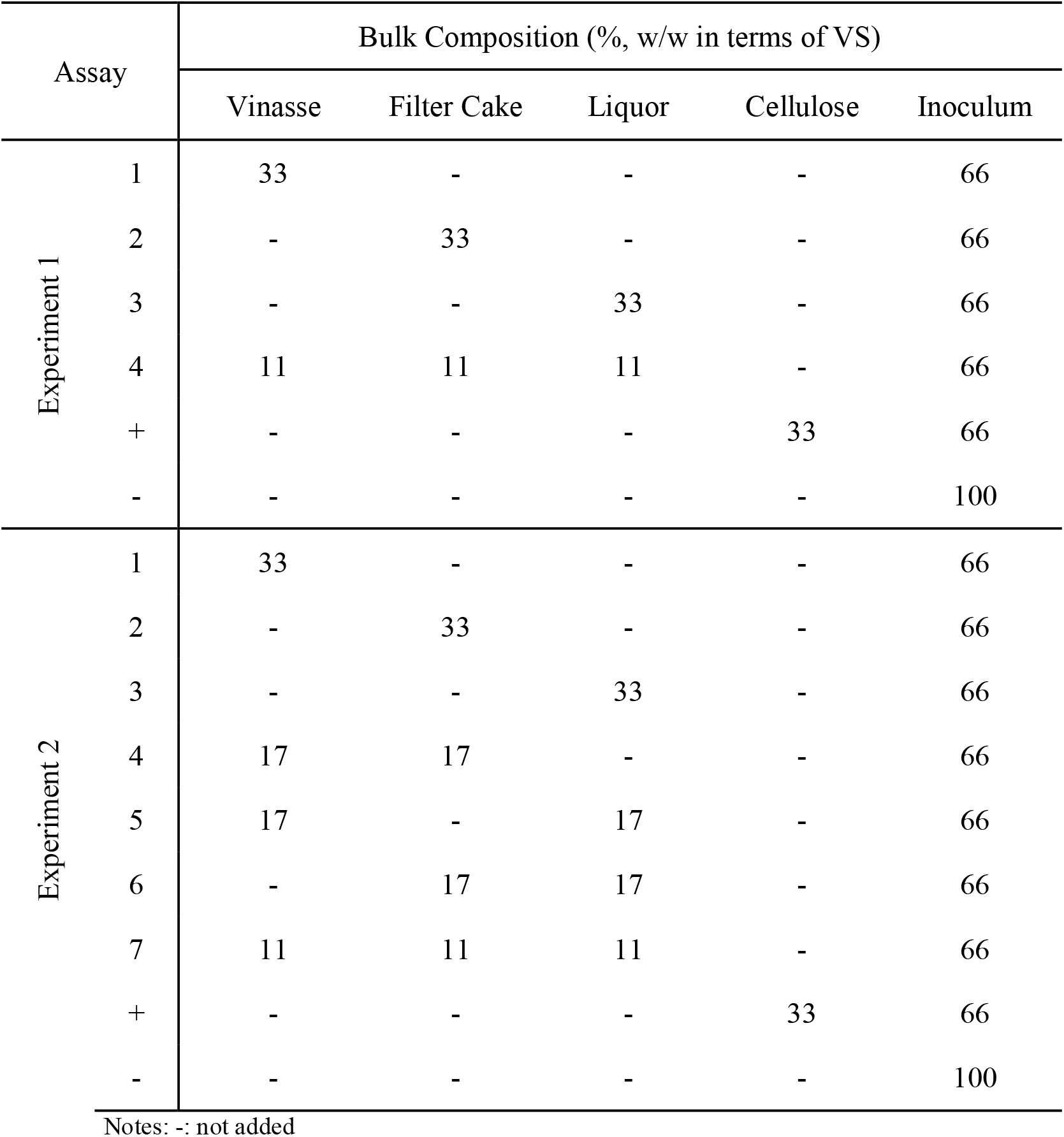
Experimental Biochemical Methane Potential (BMP) design for this study.

### 2.4. Kinetics

Co-digestion usually shows a multiple successive CH_4_ production arrangement due to the difference between the biodegradability of each substrate. It is expected that the mixture of substrates leads to a better overall performance than co-digesting each substrate separately. A modified stacked sigmoidal function (Equation 4), based on Boltzmann double sigmoid [27], was used for modeling CH_4_ volumetric production in time. The mathematical adjustment was proposed in the present work to better predict the behavior of the residues about microbial community in the production of biogas since a co-digestion process was carried out and a residue that has not been reported in the literature is being used AD systems. Based on the parameters that the model provided, it was possible to obtain better conclusions regarding the BMP of the residues.

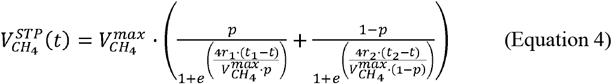

Where 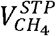 is the specific CH_4_ production in time (NmLCH_4_ g VS^-1^), 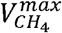 is the maximum specific volumetric production reached in the experiment (NmLCH_4_ g VS^-1^), *p* is the proportion between ordinates values of 1^st^ and 2^nd^ stacked sigmoid, *t_1_* and *t_2_* are the time which production of the 1^st^ and 2^nd^ sigmoidal pattern reaches the maximum rate (d), *r_1_* and *r_2_* are the maximum rate of CH_4_ production for the 1^st^ and 2^nd^ sigmoidal pattern, respectively (NmLCH_4_ gVS^-1^ d^-1^).

The parameter 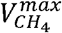 could be considered as a BMP of the assay since it represents the asymptotic maximum production of CH_4_ (when 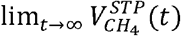). The main advantage of adopting 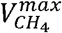 as the BMP is that it takes into consideration the trending of all experimental data, especially those in the step formed in the end phase of each batch. Thus, this approach to calculate the BMP is more precise than using the average of the last values of specific accumulated CH_4_ production.

This model assesses the maximum rate for CH_4_ production directly through both *r_1_* and *r_2_* parameters. Different from a classical Boltzmann sigmoidal function, all parameters in this model have a physical meaning. Thus, they could be useful to evaluate the studied process and for further scale-up work based on this present research. All data were processed and fitted using the software Microcal Origin^©^ 2016.

### 2.5 Physicochemical analysis

#### 2.5.1 Biogas composition

Gas chromatography (Construmaq MOD. U-13, São Carlos) analyses were performed to measure the concentration of CH_4_. The carrier gas was hydrogen (H_2_) gas (30 cm s^-1^) and the injection volume was 3 mL. The stationary phase was a 3-meter-long stainless-steel packed column (Bio-rad HPX-87H), a diameter of 1/8” with a Molecular Tamper 5A for separation of O_2_, N_2_, and CH_4_. Detection was performed through a thermal conductivity detector (TCD). Equipment was equipped with a specific injector for CH_4_, with a temperature of 350°C, an external stainless-steel wall, and an internal refractory ceramic wall. The detection limit for CH_4_ was 0.1 ppm.

#### 2.5.2 Organic Matter

The organic matter content of samples was determined in triplicate according to the Standard Methods for the Examination of Water and Wastewater [28] by the 5220B method for COD determination (digestion and spectrophotometry) and 2540 method for the solid series characterization. The solid series methodology accounted for the concentration of total (TS), and VS solids in the residues characterization.

#### 2.5.3 Sugars and acids

Concentrations of sugars and organic acids were determined in triplicate by High-Performance Liquid Chromatography (HPLC, Shimadzu^®^), composed by pump equipped apparatus (LC-10ADVP), automatic sampler (SIL-20A HT), a CTO-20A column at 43°C, (SDP-M10 AVP), Aminex HPX-87H column (300 mm, 7.8 mm, BioRad) and a refractive index detector. The mobile phase was H_2_SO_4_ (0.01 N) at 0.5 ml min^-1^.

Furfural and HMF were quantified using a Hewlett-Packard RP-18 column and acetonitrile-water (1:8 vv^-1^) containing 1% (ww^-1^) acetic acid as eluent in a flow rate of 0.8 mL min^-1^ and a UV detector at 274 nm.

#### 2.5.4 Macro and micronutrient and elementary analysis

Elementary analysis and macro and micronutrient analyses were performed at the Biomass Characterization and Analytical Calibration Resources Laboratory (LRAC), Unicamp. To determine the micronutrients, the substrate samples’ ashes were analyzed using the X-ray fluorescence equipment (brand: Panalytical, model: Axios 1KW). The ashes were prepared as is described in Standard Methods for the Examination of Water and Wastewater [28] for solid series analysis (2540 method). The elementary analysis was possible only for solid samples, i.e., filter cake, by using an elementary carbon, nitrogen, hydrogen, and sulfur analyzer (Brand: Elementar, Model: Vario MACRO Cube; Hanau, Germany).

#### 2.5.5 Total lignin (phenolic compounds)

Total lignin (soluble + insoluble lignin) content in deacetylation liquor was determined according to [29]. Acid hydrolysis was performed in pressure glass tubes with H_2_SO_4_ at 4% (w/w) final concentration and autoclaved at 121°C for 1 h. The resulting suspension was filtered and the filtrate was characterized by chromatography to determine concentrations of furan aldehydes (furfural and hydroxymethylfurfural (HMF) – as described in Section 2.5.3).

Insoluble lignin was gravimetrically determined as the solid residue from hydrolysis. For the soluble lignin, an aliquot of the hydrolysate obtained in the acid hydrolysis step was transferred to a flask with distilled water and the final pH was adjusted to 12 with a solution of 6.5 mol L^-1^ NaOH. Soluble lignin was determined from UV absorption at 280 nm using Equation 5.

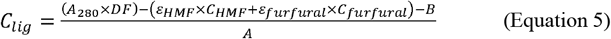

Where C_lig_ is the soluble lignin concentration in hydrolysate (g L^-1^), A280 is the absorbance of hydrolysate at 280 nm, DF is the dilution factor, ε_HMF_ is the absorptivity of HMF (114.00 L g^-1^cm^-1^ – experimental value), ε_furfural_ is the absorptivity of furfural (146.85 L g^-1^cm^-1^ – experimental value), C_HMF_ is the HMF concentration in hydrolysate (g L^-1^), C_furfural_ is the furfural concentration in hydrolysate (g L^-1^), B is the linear coefficient (0.018 – experimental value), and A is the angular coefficient equal to absorptivity of lignin (23.7 L g^-1^cm^-1^ – experimental value).

## 3. RESULTS AND DISCUSSION

### 3.1 Characterization of substrates

Table 2 shows the general characterization of substrates and inoculum. The COD value of vinasse was within the wide range generally found in the literature (15-35 g O_2_ L^-1^) [7, 8], as well as the VS content (0.015-0.020 g mL^-1^) [9], while TS content was slightly higher than previously reported (0.020-0.024 g mL^-1^) [8]. For the filter cake, the TS value was higher than normally reported (literature:0.21-0.28 g mL^-1^) [3], while VS content was much lower (literature:0.70-0.74 g mL^-1^) [13]. Such variations reflect the variability of ethanol production processes and the agricultural procedures affecting biomass characteristics, as well as the seasonality of sugarcane, already stated [8].

**Table 2.** Main parameter characterization for different substrates and inoculum.

| Residue | COD<br>(gO<sub>2</sub> L<sup>-1</sup>) | Volatile Solids<br>(g mL<sup>-1</sup>) | Total Solids<br>(g mL<sup>-1</sup>) | Fixed total Solids<br>(g mL<sup>-1</sup>) | pH | Total Lignin<br>(phenolic<br>compounds) (g L<sup>-1</sup>) |
| --- | --- | --- | --- | --- | --- | --- |
| Vinasse <sup>a</sup> | 28.81 ± 0.91 | 0.0184 ± 0.0002 | 0.0260 ± 0.0063 | 0.0077 ± 0.0005 | 4.50 ± 0.35 | -- |
| Filter Cake <sup>a</sup> | -- | 0.2021 ± 0.0005 | 0.3173 ± 0.0009 | 0.1152 ± 0.0004 | -- | -- |
| Deacetylation liquor <sup>a</sup> | 32.90 ± 0.27 | 0.0163 ± 0.0006 | 0.0215 ± 0.0021 | 0.0112 ± 0.0001 | 12.40 ± 0.13 | 5.50 |
| Iracema Mill Inoculum <sup>a</sup> | 12.70 ± 0.42 | 0.0076 ± 0.0019 | 0.0154 ± 0.0003 | 0.0078 ± 0.0000 | 7.45 ± 0.58 | -- |
| Slaughterhouse Inoculum <sup>a</sup> | 20.01 ± 0.78 | 0.0466 ± 0.0076 | 0.0547 ± 0.0001 | 0.0081 ± 0.0000 | 7.32 ± 0.27 | -- |

<sup>a</sup>Three replicates average ± standard deviation; --:not determined

Elementary characterization of filter cake showed that it is mainly composed of 0.16% sulfur, 1.73% nitrogen, 31.56% carbon, and 3.11% hydrogen (in %TS). The values for S and N are close to those found in the literature (0.18% and 1.76%, respectively) [30]; however, the C value is below what is normally reported (40-42%) [18]. It resulted in the C:N ratio of the filter cake of 18:1, below what is recommended for AD, which is 20-40: 1 [31].

Slaughterhouse inoculum presented higher values of COD, VS, and TS than the inoculum of the sugarcane mill (Table 2), already predicting that it may have a better development for biogas production as it probably contains high cellular mass, i.e. microbiological content. Additionally, the slaughterhouse inoculum visually presented a good quality granular appearance from UASB reactors, while the mill inoculum had a liquid aspect. Both pHs were neutral, as expected for anaerobic inocula.

The deacetylation liquor presented a strong alkali characteristic since it came from a mild alkaline pretreatment of sugarcane straw to remove acetyl groups and promote lignin solubilization [12]. Alkaline pretreatment is typically used in lignocellulosic materials such as wheat straw and sugarcane bagasse, thus decreasing its recalcitrance [4]. According to the deacetylation liquor composition (Table 2 and Table 3), a large amount of lignin fractions was detected (phenolic compounds) and high amounts of acids that can be transformed into CH_4_, thus showing evidence of a potential high experimental CH_4_ production. Several types of pre-treatments are currently carried out with sugarcane lignocellulosic materials, such as chemical (acid, alkaline), biological, physical, and physicochemical, in which different types of residues are generated with different characteristics, pH, carbohydrate composition, and lignin content [32]. Thus, it is difficult to make comparisons with the literature. It is worth mentioning that the deacetylation liquor obtained from this work could be specially benefitted for the co-digestion with vinasse due to its basic character. The deacetylation liquor could neutralize the low pH of vinasse without adding large amounts of an alkalizing agent, proving some possible economic benefits of the AD system. The need to alkalize vinasse before AD is an economic disadvantage in terms of implementing this process in sugarcane mills [33]. The presence of C6 and C5 sugars, such as glucose, xylose, arabinose, and the presence of oligosaccharides, such as arabinoxylan and glucan (Table 3), is also highlighted which can be used by the anaerobic microbial community for conversion to CH_4_, although constraints of AD from C5 sugars are commonly reported [34, 35].

**Table 3.** Acid and sugar content of liquid substrates

|  | Vinasse<sup>a</sup> (mg L<sup>-1</sup>) | Deacetylation liquor<sup>a</sup> (mg L<sup>-1</sup>) |
| --- | --- | --- |
| Acetate | 2215.91 ± 0.80 | 3670.00 ± 0.89 |
| Isobutyrate | 2076.27 ± 1.50 | 0.00 |
| Formate | 0.00 | 63.00 ± 1.35 |
| Malate | 4944.00 ± 0.48 | 0.00 |
| Lactate | 2618.17 ± 0.98 | 0.00 |
| Glucose | 0.00 | 85.204 ± 2.45 |
| Glucan | -- | 626.00 ± 1.12 |
| Fructose | 1045.25 ± 0.43 | 0.00 |
| Arabinose | -- | 26.00 ± 0.44 |
| Xylose | -- | 35.00 ± 0.95 |
| Arabinoxylan | -- | 1747.00 ± 2.32 |

<sup>a</sup>Mean of three replicates ± standard deviation; --: not determined

High values of acetic acid were obtained for both vinasse deacetylation liquor (Table 3), in which this volatile fatty acid was reported as important and essential for the acetotrophic methanogenic metabolic route [36]. Also, Wang et al. [37] noted that concentrations of acetic acid and butyric acid of 2400 and 1800 mg L^-1^, respectively, did not result in significant inhibition of methanogenic activity. Lactic acid was found in high concentrations in vinasse, and it is usually degraded to propionic acid, which is an undesirable terminal fermentation product; thus high concentrations of propionic acid can result in methanogenesis failure [37]. Moreover, the high concentration of lactic acid in vinasse may result in inhibitory effects for CH_4_ production, highlighting the potential advantage of applying the co-digestion to balance the volatile fatty acid composition in the medium. Vinasse also presented malic acid which is generally from the sugarcane plant [37] and isobutyric acid, contributing to its acidic pH.

Table 4 shows the macro and micronutrient concentrations detected in the substrates. As no external micronutrient solution was added to the experiments, the effects of the nutrient content of the residues could be ascertained by comparing their BMP behavior with the positive control test (cellulose), which had an absence of nutrients. Menon et al. [38] showed optimal concentrations of 303 mg L^-1^ Ca, 777 mg L^-1^ Mg, 7 mg L^-1^ Co, and 3 mg L^-1^ Ni that increased biogas productivity by 50% and significantly reduced the processing time. Filter cake presented higher concentrations of the aforementioned micronutrients, except for Ni which was not detected. It is known that an excess of these compounds may cause inhibitory effects on AD, increasing the lag phase of the process [39] or reducing the specific CH_4_ production [40]. A considerable amount of S was also detected in filter cake, which could decrease CH_4_ formation from acetate due to the sulfate-reducing bacteria activity. Such bacteria compete by using acetate for sulfide production and can even inhibit methanogenesis activity, leading the process to failure [41]. Al and Fe were also present in inhibitory concentrations, which were reported in the literature with values greater than 2.5 g L^-1^ and 5.7 g L^-1^, respectively [42]. Mg and Ca concentrations were also much above what is recommended for AD (ideally around 0.02 mg L^-1^ and 0.03 mg L^-1^, respectively), which may also contribute to the inhibition of the process [43]. High concentrations of Mg ions stimulate the production of single cells of microorganisms with high sensitivity for lysis, leading to a loss of acetoclastic activity in anaerobic reactors, while high Ca concentrations can lead to an accumulation of biofilm, which impairs methanogenic activity and may also cause buffering capacity loss of the essential nutrients for AD [42]. On the other hand, cobalt (Co) was detected only in this substrate, within the stimulating concentration range for methanogenesis [44]. These findings reinforce the need of using co-substrates to dilute the potential inhibitory effects caused by excessive concentrations of nutrients in the filter cake while taking advantage of beneficial effects that certain components of its composition may provide.

**Table 4.** Macro and micronutrient concentration of substrates and inocula

| Nutrients | Vinasse<br>(g L<sup>-1</sup>TS<sup>-1</sup>) | Filter Cake<br>(g L<sup>-1</sup>TS<sup>-1</sup>) | Deacetylation<br>liquor<br>(g L<sup>-1</sup>TS<sup>-1</sup>) | Slaughterhouse<br>Inoculum<br>(g L<sup>-1</sup>TS<sup>-1</sup>) | Iracema Mill<br>Inoculum<br>(g L<sup>-1</sup>TS<sup>-1</sup>) |
| --- | --- | --- | --- | --- | --- |
| Al | 0.0137 | 16.1825 | 0.4164 | 0.3719 | 0.0402 |
| Ba | 0.0000 | 0.0000 | 0.0000 | 0.0014 | 0.0000 |
| Br | 0.0024 | 0.0000 | 0.0000 | 0.0003 | 0.0013 |
| Ca | 0.4682 | 6.0729 | 0.4055 | 0.2349 | 0.7875 |
| Cl | 1.2856 | 0.0657 | 0.2546 | 0.0572 | 0.7764 |
| Co | 0.0000 | 0.0330 | 0.0000 | 0.0023 | 0.0000 |
| Cr | 0.0000 | 0.0252 | 0.0000 | 0.0030 | 0.0000 |
| Cu | 0.0000 | 0.0230 | 0.0031 | 0.1097 | 0.0016 |
| Fe | 0.0163 | 10.6633 | 0.3227 | 1.1316 | 0.1062 |
| Ga | 0.0000 | 0.0051 | 0.0008 | 0.0003 | 0.0000 |
| Ge | 0.0000 | 0.0000 | 0.0021 | 0.0008 | 0.0000 |
| K | 2.6078 | 0.9487 | 1.1680 | 0.2709 | 1.7843 |
| Mg | 0.5372 | 1.4284 | 0.1286 | 0.1155 | 0.3595 |
| Mn | 0.0047 | 0.2864 | 0.0123 | 0.0073 | 0.0103 |
| Mo | 0.0000 | 0.0000 | 0.0000 | 0.0063 | 0.0004 |
| Na | 0.0849 | 0.0000 | 10.4902 | 0.7204 | 0.0000 |
| Nb | 0.0000 | 0.0040 | 0.0000 | 0.0000 | 0.0000 |
| Ni | 0.0000 | 0.0000 | 0.0000 | 0.0028 | 0.0000 |
| P | 0.0913 | 2.9929 | 0.1120 | 0.5496 | 0.2029 |
| Pb | 0.0000 | 0.0086 | 0.0000 | 0.0015 | 0.0000 |
| Rb | 0.0039 | 0.0042 | 0.0000 | 0.0006 | 0.0030 |
| Si | 0.5384 | 0.5304 | 1.1620 | 0.4663 | 0.0891 |
| S | 0.0739 | 18.1076 | 0.3345 | 0.5495 | 0.3779 |
| Sr | 0.0021 | 0.0419 | 0.0025 | 0.0025 | 0.0033 |
| Ti | 0.0013 | 1.9849 | 0.0374 | 0.0200 | 0.0025 |
| V | 0.0000 | 0.0529 | 0.0000 | 0.0000 | 0.0000 |
| W | 0.0000 | 0.0000 | 0.0000 | 0.0033 | 0.0000 |
| Zn | 0.0006 | 0.0491 | 0.0043 | 0.1292 | 0.0163 |
| Zr | 0.0000 | 0.0537 | 0.0000 | 0.0010 | 0.0000 |

Deacetylation liquor presented the main micronutrients in milder concentrations considered important for the development of methanogenic *Archaea*, such as Fe, Zn, Cu, Mn, which stimulate reactions catalyzed by metalloenzymes, the formation of cytochromes, and ferroxins [45]. However, high concentrations of Si and especially Na were detected. The presence of large amounts of Si is intrinsic to lignocellulosic materials [46]. The use of Si as a trace element for AD is rarely reported since it is often either volatilized in the biogas produced or else it remains in the digested material [47], not affecting the AD process. The Na can cause an inhibitory effect on the methanization of volatile fatty acids (mainly propionic acid) in concentrations between 3 to 16 g L^-1^; however, for glucose-rich-substrates, this Na concentration does not significantly affect methanogenesis [48]. Methanogenic archaea can also adapt to high Na concentration, leading to high CH_4_ conversions [48]. Vinasse did not present known inhibitory concentrations for the assessed macro and micronutrients [42].

Comparing the nutritional content of the inocula, the slaughterhouse inoculum presented a wider range of components in mild concentrations, indicating richer anaerobic microbial activity than the inoculum from the sugarcane mill, especially Co, Ni, Fe content that together allows better development of methanogenic activity [49]. The mill’s inoculum, on the other hand, had neither Co nor Ni trace metals and much lower Fe concentration. The nutritional poverty of the latter inoculum is accompanied by high K content, consistent with the vinasse treatment, a K-rich substrate.

### 3.2 BMP: Experiment 1

The main results of the BMP tests of Experiment 1 are presented in Table 5, including the experimental values and the kinetic parameters obtained from the mathematical modeling and Tukey anlysis. The respective curves of the cumulative volume of produced CH_4_ are presented in Figure 1. Co-digestion of substrates enhanced CH_4_ production when compared to the AD of isolated substrates. However, the positive control (cellulose) did not reach the minimum recommendable BMP value (352 NLCH_4_ kgVS^-1^) to validate results as maximum potential values for specific CH_4_ production [50]. That indicates that the maximum capacity for producing CH_4_ from the assessed substrates may not have been reached. The most probable cause for this lack of performance observed in Experiment 1 might be occurred due to the inoculum. Although cellulose digestibility was low, high digestibilities were obtained for liquid substrates (vinasse and deacetylation liquor), which indicates that the presence of nutrients in the substrates (Table 4) has positively affected the inoculum activity as no nutritional supplementation was added in all assays. According to Menon et al. [38], the use of micronutrients remedies AD with a focus on CH_4_ production in thermophilic process and increases biogas productivity. Also, a high concentration of acetate in vinasse and deacetylation liquor could be an important factor to boost CH_4_ production, since acetate is the sole substrate used by acetoclastic methanogens *Archaea*. Despite the high organic content, the filter cake showed low biodigestibility compared to the other residues (53%). It is likely the excess of micronutrients and S concentrations negatively affected methanogenesis (Table 4). Besides, the physical limitations on the biological process due to the higher TS content (at least 12-fold greater than the other co-substrates) (Table 2) was another major limiting factor. The absence of stirring may have hindered the mass transfer between the substrate and the inoculum, reducing the microbiological reactions involved in the AD process and not allowing to achieve higher BMP values [51].

**Table 5.** Values of experimental BMP, biodigestibility, pH (initial and final), and kinetic parameters of isolated and co-digested substrates of Experiment 1.

| Parameters | Cellulose | Vinasse<sup>a</sup> | Deacetylation Liquor<sup>b</sup> | Filter Cake<sup>c</sup> | (Vinasse + Liquor + Filter Cake)<sup>b</sup> |
| --- | --- | --- | --- | --- | --- |
| Values of experimental BMP |  |  |  |  |  |
| TBMP (NmLCH<sub>4</sub> gVS<sup>-1</sup>) | 415 | 548 | 706 | 900 | -- |
| <sup>1</sup>BMP (NmLCH<sub>4</sub> gVS<sup>-1</sup>) | 282 ± 13 | 476 ± 13 | 610 ± 50 | 353 ± 38 | 660 ± 49 |
| Biodigestibility (%) |  |  |  |  |  |
| <sup>2</sup>Average | 68 | 87 | 46 | 40 | -- |
| <sup>3</sup>Model | 71 | 89 | 95 | 45 | -- |
| pH initial | 7.98 ± 0.47 | 7.13 ± 0.02 | 7.90 ± 0.85 | 7.77. ± 0.25 | 7.91 ± 0.01 |
| pH final | 7.69 ± 0.75 | 7.81 ± 0.15 | 7.80 ± 0.08 | 7.83 ± 0.01 | 7.78 ± 0.47 |
| Kinetic Model Parameters |  |  |  |  |  |
| $V_{CH_4}^{max}$ (NmLCH<sub>4</sub> gVS<sup>-1</sup>) | 295.83* | 487.27* | 673 ± 20 | 404.69* | 688 ± 9 |
| $p_1$ | 0.39 ± 0.02 | 0.48 ± 0.01 | -- | 0.53 ± 0.02 | -- |
| $r_1^{max}$ (NmLCH<sub>4</sub> gVS<sup>-1</sup> d<sup>-1</sup>) | 3.6 ± 0.06 | 7.4 ± 0.5 | 16 ± 0 | 5.5 ± 0.4 | 12 ± 0 |
| $t_1$ (d) | 25 ± 2 | 11 ± 1 | 57 ± 1 | 29 ± 2 | 32 ± 1 |
| $r_2^{max}$ (NmLCH<sub>4</sub> gVS<sup>-1</sup> d<sup>-1</sup>) | 15 ± 3 | 13 ± 1 | -- | 11 ± 2 | -- |
| $t_2$ (d) | 75 ± 1 | 72 ± 0 | -- | 76 ± 1 | -- |
| $R^2$ | 0.89 | 0.97 | 0.99 | 0.96 | 0.99 |

Notes:<sup>1</sup>Average value of the replicates following the equation $3 \pm$ standard deviation <sup>2</sup>Calculated considering BMP; <sup>3</sup>Calculated considering $V_{CH_4}^{max}$ , --: not determined,
\*Parameters values forced. Values within a row with the same letter are not significantly different at 5% probability by Tukey

**Fig. 1.**
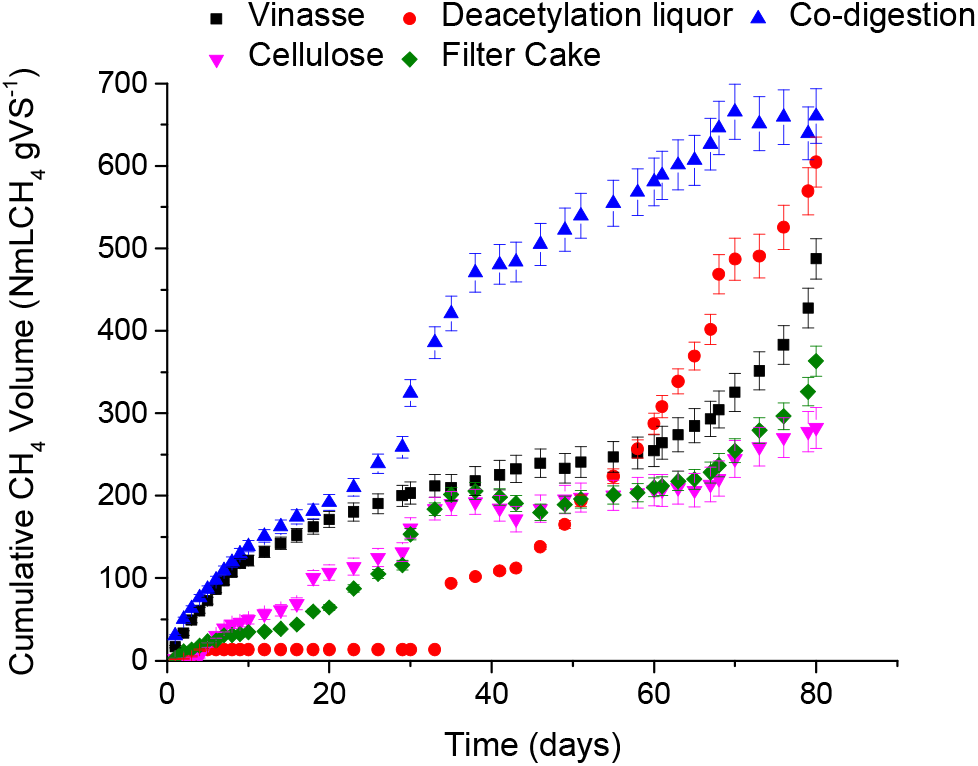
Cumulative methane volume from BMP of Experiment 1

The pH of the assay was adjusted to between 7-8 at the beginning of the experiment and throughout the experiment it remained in this range, occurring neither acidification nor alkalinization.

Despite the high BMP value and high digestibility of deacetylation liquor, its lag phase was significantly long: CH_4_ was produced only after 40 days. The long lag phase can be caused by the presence of pre-treatment inhibitors for alcoholic microorganisms, which are commonly reported [52, 53]. However, the presence of furfural or HMF, commonly reported as inhibitors, was not identified. This fact raises two hypotheses: an excess of Na, which may have led to a longer time for methanogenic community adaptation (Section 3.1); the presence of fractions of lignin and derived compounds, which may have caused the observed “delay” in the release of organic matter in the environment to access the microbiota. The degradation process of lignin to be used in AD is quite complex, in which some steps are involved before the acetogenesis process [54]. The lignin polymer is first depolymerized and then solubilized, in which different lignin monomers are formed, with varying chain sizes, such as phenylpropanoid derivates with a carboxylic acid, alcohol, or amine groups. After this stage, these monomers undergo a wide variety of peripheral pathways to form other intermediates, which are the central monoaromatic intermediate, such as resorcinol (trihydroxibenezene). These elements proceed to the dearomatization and cleavage stage of the aromatic ring, forming aliphatic acids. This aliphatic acids enter in acidogenesis phase and they are degraded into volatile fatty acids to continue in the following AD stages [55]. Thus, the long lag phase of deacetylation liquor AD observed in the BMP test may have happened due to the long process of degradation of lignin fractions and derived compounds, since lignin fractions (i.e., phenolic compounds) were detected in this substrate at significant levels (Table 3).

The biodigestibility predicted by the kinetic modeling showed higher values (from 2% to 10% higher) than the ones calculated from the experimental BMP, as a trend to produce CH_4_ after ending the experiments was detected by the fitted model. This behavior indicates a trend towards bigger accumulated production of biogas in a longer time than the experiments were conducted. This fact was especially observed for the deacetylation liquor: the model predicted a larger lag phase (57 *vs* 34 days) and BMP (673 *vs* 610 NmLCH_4_ gVS^-1^) values than the observed experimentally. These data confirm that the presence of phenolic compounds (lignin and derivatives) may have caused this long time in the lag phase.

The fitting of the kinetic model was not adequate for any of the studied substrates in Experiment 1. As depicted in Table 5, cellulose, vinasse, and filter cake must have the 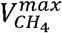 value forced to the average of the CH_4_ accumulated production at the end of each respective assay so the model could fit. The adjusted curves are presented in the Supplementary Materials (Figure 1SM). It is noteworthy that although the experiments were completed according to VDI 4630 [24] (the process was finished when production of biogas per batch was less than 1% of the accumulated gas production), the kinetic model showed a trend of increase in CH_4_ production over a longer period. For this reason, the values of 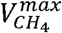 can be considered the maximum BMP. Despite this fact, no distinguishable step was observed in Figure 1, indicating that those assays might have reached the stationary phase. The absence of this last step in the experimental data leads to a lack of goodness of fitting, although reaching a good coefficient of correlation (R^2^).

Experiment 1 showed that deacetylation liquor did not show a double sigmoid pattern, indicating that all the CH_4_ production occurred in one single step. It confirms the peculiar behavior of this residue from producing CH_4_: despite the large lag phase (t1 = 57 days) and the delay in the organic matter degradation, the overall rate for CH_4_ conversion (observed by the 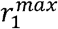 value) was the highest among the substrates, which occurred in one phase. This same pattern could be observed in the co-digestion of vinasse, liquor, and filter cake altogether, suggesting that the deacetylation liquor could have improved the process of CH_4_ production. Another strong evidence that the deacetylation liquor is the substrate responsible for improving the co-digestion is the high value of the apparent kinetic parameter 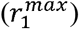 in this experiment after the experiment with deacetylation liquor. Co-digestion increased the CH_4_ production rate by 38% and 54% when compared to the isolated AD from vinasse and filter cake, respectively. This behavior is also confirmed by the analysis of ANOVA and the Tukey test that showed a significant difference between the treatments of the residues alone, however, concerning the deacetylation liquor and the co-digestion, there was no significant difference, indicating that the deacetylation liquor was responsible for the increase in CH_4_ production in co-digestion.

### 3.3 BMP: Experiment 2

Table 6 shows the main results from the BMP tests of Experiment 2, including the experimental values, the kinetic parameters obtained from the mathematical modeling and Tukey analysis. The adjusted curves for this experiment can be consulted in the Supplementary Materials (Figure 2SM). Unlike Experiment 1, high biodigestibility of cellulose (positive control) was reached (> 85%), thus validating the BMP tests as maximum experimental CH_4_ production from assessed substrates [50]. The BMP values obtained in Experiment 2 are, thus, the representative ones for the assessed residues. This fact indicates better quality of anaerobic inoculum from the poultry slaughterhouse treatment when compared to the inoculum from sugarcane vinasse treatment. Biogas production constraints from vinasse on a scale (e.g. variation of vinasse composition throughout the season, AD reactor shutdown in the vinasse offseason) reflects the lack of robustness of the inoculum due to its continuous need for adaptation to the substrate, which weakens the microbial activity. It is noteworthy that the kinetic model prediction was close to the experimental values of BMP, reinforcing that the maximum BMP was experimentally reached: the biodigestibility (Table 6) calculated from both methods varied only by 3% on average. It confirms the robustness of the inoculum and proves the representativity of BMP results.

**Fig. 2.**
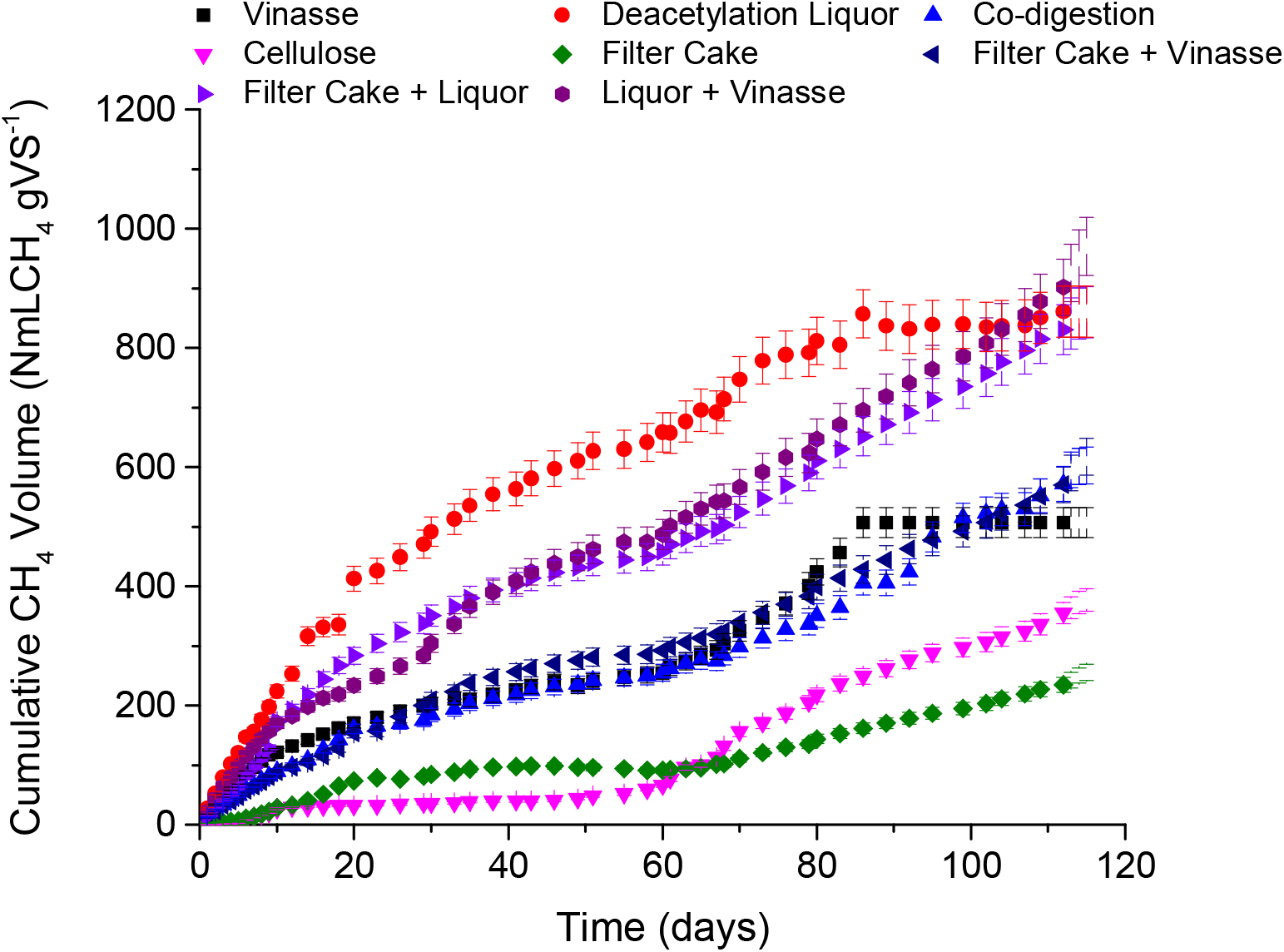
Cumulative methane volume from BMP of Experiment 2

**Table 6.** Values of experimental BMP, biodigestibility, pH (initial and final), and kinetic parameters of isolated and co-digested substrates of Experiment 2.

| Parameters | Cellulose | Vinasse <sup>a</sup> | Deacetylation<br>Liquor <sup>b</sup> | Filter<br>Cake <sup>c</sup> | (Vinasse + Liquor<br>+ Filter Cake) <sup>a</sup> | (Vinasse +<br>Filter Cake) <sup>a</sup> | (Vinasse +<br>Deacetylation<br>Liquor) <sup>b</sup> | (Deacetylation<br>Liquor + Filter<br>Cake) <sup>b</sup> |
| --- | --- | --- | --- | --- | --- | --- | --- | --- |
| Values of experimental BMP |  |  |  |  |  |  |  |  |
| TBMP (NmLCH<sub>4</sub><br>gVS<sup>-1</sup>) | 415 | 548 | 706 | 900 | -- | -- | -- | -- |
| <sup>1</sup>BMP (NmLCH<sub>4</sub><br>gVS<sup>-1</sup>) | 380 ± 7 | 507 ± 6 | 853 ± 43 | 262 ± 2 | 605 ± 88 | 614 ± 8 | 971 ± 72 | 861 ± 24 |
| Biodigestibility (%) |  |  |  |  |  |  |  |  |
| <sup>2</sup>Average | 92 | 93 | 121 | 29 | -- | -- | -- | -- |
| <sup>3</sup>Model | 89 | 94 | 122 | 30 | -- | -- | -- | -- |
| pH initial | 7.52 ± 0.45 | 7.47 ± 0.02 | 7.79 ± 0.15 | 7.59 ±<br>0.89 | 7.51 ± 2.85 | 7.45 ± 1.45 | 7.63 ± 0.39 | 7.71 ± 0.42 |
| pH final | 7.38 ± 0.08 | 7.12 ± 0.76 | 7.89 ± 0.61 | 7.49 ±<br>0.73 | 7.35 ± 0.92 | 7.32 ± 0.56 | 7.55 ± 0.03 | 7.87 ± 0.25 |
| Kinetic Model Parameters |  |  |  |  |  |  |  |  |
| $V_{CH_4}^{max}$ (NmLCH<sub>4</sub><br>gVS<sup>-1</sup>) | 368 ± 7 | 513 ± 3 | 863 ± 9 | 274 ± 9 | 717 ± 119 | 797 ± 110 | 1298 ± 660 | 1021 ± 70 |
| $p_1$ | 0.07 ± 0.01 | 0.44 ± 0.01 | 0.58 ± 0.03 | 0.32 ±<br>0.02 | 0.27 ± 0.08 | 0.28 ± 0.07 | 0.27 ± 0.32 | 0.31 ± 0.04 |
| $r_1^{max}$ (NmLCH<sub>4</sub><br>gVS<sup>-1</sup> d<sup>-1</sup>) | 5.6 ± 4.6 | 7.5 ± 0.4 | 20 ± 1 | 5.1 ± 0.4 | 7.2 ± 1.6 | 7.0 ± 0.4 | 8.3 ± 2.1 | 17 ± 1 |
| $t_1$ (d) | 7.4 ± 1.1 | 11 ± 1 | 12 ± 1 | 14 ± 1 | 13 ± 2 | 17 ± 1 | 18 ± 3 | 12 ± 0 |
| $r_2^{max}$ (NmLCH<sub>4</sub><br>gVS<sup>-1</sup> d<sup>-1</sup>) | 7.3 ± 0.3 | 11 ± 1 | 7.3 ± 0.6 | 3.7 ± 0.1 | 7.7 ± 0.7 | 6.4 ± 0.2 | 9.1 ± 0.8 | 7.9 ± 0.3 |
| $t_2$ (d) | 79 ± 1 | 75 ± 0 | 62 ± 2 | 93 ± 2 | 94 ± 8 | 100 ± 7 | 102 ± 25 | 89 ± 4 |
| $R^2$ | 0.97 | 0.99 | 0.98 | 0.99 | 0.90 | 0.99 | 0.95 | 0.99 |

Notes: <sup>1</sup>Average value of the replicates following the equation $3 \pm$ standard deviation; <sup>2</sup>Calculated considering BMP; <sup>3</sup>Calculated considering $V_{CH_4}^{max}$ , --: not determined. Values within a row with the same letter are not significantly different at 5% probability by Tukey

Lower filter cake BMP was obtained when compared to Experiment 1. The physical characteristics of inocula could have played a role in this case: the inoculum from poultry slaughterhouse treatment was composed of very well-formed granules (traditional Upflow Anaerobic Sludge Blanket - UASB sludge), while inoculum from vinasse treatment was liquid without any granules. The mass transfer resistance in anaerobic granules might limit CH_4_ production, since the larger the granule, the greater the resistance to mass transfer [56], which may have been attenuated with the liquid inoculum for the filter cake access. Additionally, in the co-digestion BMP tests, the highest value of BMP was obtained with only liquid substrates (deacetylation liquor + vinasse) while using filter cake as co-substrate caused a decrease in BMP values (Table 6). It reinforces that the mass transfer phenomena have an important influence on CH_4_ production from filter cake, which must be considered for a reactor operation and inoculum sludge choice. The excess concentrations of some macro and micronutrients already discussed (Section 3.1) may also have contributed to the lower BMP.

Experimental BMP of deacetylation liquor showed an atypical result, as it was higher than its TBMP value. Deacetylation pretreatment liquor (with the alkaline character) has favorable characteristics for CH_4_ production because it reduces the degree of inhibition on CH_4_ fermentation [57], which may explain its high BMP value (Table 6). However, the lower TBMP than BMP implies the possibility that all organic matter in the deacetylation liquor was not accounted for in the COD value, underestimating the value of TBMP. Remnants of insoluble lignin may not have been quantified in the COD analysis [58] and during the BMP tests, they may have been hydrolyzed and made available as soluble lignin [55, 58]. CH_4_ production from soluble lignin was already reported [59]. It is also worth mentioning that trace metals can act as catalysts favoring the depolarization of the soluble lignin in the liquid medium, thus leaving more organic matter available [60]. The inoculum used in Experiment 1 had lower metal content when compared to the slaughterhouse inoculum of Experiment 2, (especially Al, Co, Fe, Cu) corroborating the hypothesis that the presence of metals may have contributed to the depolarization of soluble lignin in the deacetylation liquor. Thus, larger metal content in poultry inoculum may lead to larger amounts of available organic matter during the BMP test, which was not accounted for in the COD value of deacetylation liquor determined in the absence of inoculum. These assumptions highlight the need for deeper further studies on CH_4_ production from liquid lignocellulosic substrates.

As in Experiment 1, the pH of Experiment 2 remained neutral throughout the operation, with no acidification or alkalinization of the medium, and no need for initial pH correction exclusively for the co-digestion test.

The co-digestion of substrates showed higher potential for CH_4_ production than the AD of isolated residues, as in Experiment 1, except for the deacetylation liquor. However, considering the context of a sugarcane biorefinery, its most abundant residue (i.e., vinasse) must be properly managed, whereby AD is an advantageous alternative as already reported [7]. The enhancement of CH_4_ production from vinasse can be achieved by adding other residues within the biorefinery boundary as co-substrates, as proved in the current work. By predicting a co-digestion reactor operation, in which the continuous stirred tank reactor (CSTR) is the traditional one [8], the disadvantage of the filter cake by having a higher ST content could be minimized due to stirring, avoiding its sedimentation and improving the substrate-inoculum contact and, therefore, resulting in increased CH_4_ production.

Kinetic modeling performed in all assays of Experiment 2 (Table 6) showed a particularly good fitting. All assays showed a clear ending step at the end of each assay, and the model represented all data without any need of forcing parameters to a value.

As showed in Experiment 1, deacetylation liquor had the highest values for both 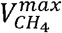 and 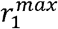 demonstrating a higher and faster CH_4_ production than any other assay. This value was 4 times greater than the 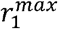 of the other isolated substrates. Codigestion of deacetylation liquor and vinasse; and deacetylation liquor and filter cake have been proved more capable of CH_4_ production than other conditions. Kinetic parameters also confirmed that co-digestion was more effective than single substrate AD for CH_4_ production, and the deacetylation liquor was probably the substrate that boosted methanogenesis in co-digestion assays. Besides, the 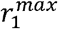 values of the deacetylation liquor were higher in the current experiment when compared to Experiment 1 (20 *vs* 16 NmLCH_4_ d^-1^). This fact confirms that in Experiment 2 the phenolic compounds (lignin and derivates) of deacetylation liquor may have been faster solubilized in shorter organic matter chains to be converted to CH_4_ (section 3.3), making its 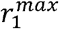 a value higher than in Experiment 1, where the lignin content took a longer period to be solubilized and thus resulting in a lower CH_4_ production rate. This corroborates that TBMP was underestimated since lignin was not fully accounted for in the COD analysis in Experiment 1. The result of the Tukey test also confirms the hypotheses raised above, since there was no significant difference at 5% probability of the BMP of deacetylation liquor, vinasse and deacetylation liquor, and filter cake and deacetylation liquor tests. This situation shows that the AD assay of the deacetylation liquor alone or of him with the other residues there will be no difference in final CH_4_ production, indicating that the deacetylation liquor was the residue that leveraged methanogenesis when co-digesting with the other two residues. Although the Tukey test did not show a significant difference between the BMP test of the vinasse, of the codigestion of the three residues, and the vinasse with the filter cake, it is notorious that the BMP values increased and a lot with the presence of the deacetylation liquor.

Figure 2 shows the curves of the cumulative volume of produced CH_4_ in Experiment 2, presenting a more accentuated behavior of AD occurring in two-phases when compared to Experiment 1: the acidogenic phase and the subsequent methanogenic phase [61]. This proves that the origin of the inoculum plays an important role in the production of CH_4_, as the same substrates were used in the two rounds of experiments. The BMP of the substrates in Experiment 2 had a shorter lag phase when compared to Experiment 1, as confirmed by the obtained kinetic parameters (t1), indicating that there was a better adaptation of the inoculum to the substrate. Gu et al. [62] observed distinct performance of biogas production using different inocula for the same substrate (rice straw), showing that some inocula were better adapted than others due to their specific enzymatic arsenal and to the degraded organic matter load capacity: the greater organic matter converted by the inoculum, the better it would be able to convert lignocellulosic residues.

The inoculum used in Experiment 2 came from a consolidated UASB reactor continuously treating poultry slaughterhouse waste, with higher organic loads fed to the reactor when compared to the inoculum used in Experiment 1 (from a reactor that has been in operation for only 4 years for the treatment of vinasse). This made the slaughterhouse inoculum more robust than mill inoculum, and, thus, more suitable and efficient to convert lignocellulosic materials, causing the smallest lag phase and making the digestion process more stable, which results in higher cumulative CH_4_ volumes [63].

## 4. CONCLUSION

Anaerobic inoculum maturity improved the slow conversion of lignin-fraction monomers into CH_4_ from deacetylation liquor. Its alkali-characteristic may contribute to the AD operational costs reduction on an industrial scale as it avoided the reactor alkalizing demand. The highest filter cake TS content indicated operational adjustments are necessary, e.g., stirring to minimize the mass transfer resistance between substrate and microorganisms. The adjusted kinetic model confirmed the maximum experimental BMP values for the robust AD inoculum. This small-scale study shows how the codigestion made use of residues positive synergisms to increase CH_4_ yield by at least 16%. This highlighted the advantage for the management of the voluminous residue of integrated 1G2G sugarcane biorefineries (vinasse) and new lignin-rich side streams derived from pretreatment technologies for sugarcane straw valorization, so far unexplored for methane production

## Supporting information

Original dataset paper

## ACKNOWLEDGEMENTS

This work was supported by São Paulo Research Foundation - FAPESP contract numbers 2018/09893-1 TO XXX2016/16438-3 to XXX, 2017/15477-8 to LBB 2015/50612-8 (FAPESP-BBSRC Thematic Project). The authors gratefully acknowledge the support of the Laboratory of Environment and Sanitation (LMAS) at the School of Agricultural Engineering (FEAGRI/UNICAMP), the National Laboratory of Biorenewables (LNBR/CNPEM), and the Interdisciplinary Center of Energy Planning (NIPE/UNICAMP).

## AUTHOR CONTRIBUTION STATEMENT

Maria Paula C. Volpi: Conceptualization, Methodology, Data curation, and Writing-original draft preparation.

Li via B. Brenelli: Methodology, Data curation, and Writing-original draft preparation.

Gustavo Mockaitis: Methodology, Data curation, and Writing-original draft preparation.

Sarita C. Rabelo: Methodology, Data curation, and Writing-original draft preparation.

Telma T. Franco: Project administration, and Funding acquisition.

Bruna S. Moraes: Conceptualization, Formal analysis, Writing - Review & Editing, Supervision.

## AVAILABILITY OF DATA AND MATERIAL

The datasets generated during and /or analyzed during the current study are available from the corresponding author on reasonable request

## CODE AVAILABILITY

Not applicable

## REFERENCES

1. Parsaee M, Kiani Deh Kiani M, Karimi K (2019) A review of biogas production from sugarcane vinasse. Biomass and Bioenergy 122:117–125. https://doi.org/10.1016/j.biombioe.2019.01.034

2. Moraes BS, Triolo JM, Lecona VP, et al (2015) Biogas production within the bioethanol production chain: Use of co-substrates for anaerobic digestion of sugar beet vinasse. Bioresour Technol 190:227–234. https://doi.org/10.1016/j.biortech.2015.04.089

3. Janke L, Leite A, Nikolausz M, et al (2015) Biogas Production from Sugarcane Waste: Assessment on Kinetic Challenges for Process Designing. Int J Mol Sci 16:20685–20703. https://doi.org/10.3390/ijms160920685

4. Rabelo SC, Carrere H, Maciel Filho R, Costa AC (2011) Production of bioethanol, methane and heat from sugarcane bagasse in a biorefinery concept. Bioresour Technol 102:7887–7895. https://doi.org/10.1016/j.biortech.2011.05.081

5. Nakanishi SC, Soares LB, Biazi LE, et al (2017) Fermentation strategy for second generation ethanol production from sugarcane bagasse hydrolyzate by Spathaspora passalidarum and Scheffersomyces stipitis. Biotechnol Bioeng 114:2211–2221. https://doi.org/10.1002/bit.26357

6. Adarme OFH, Baêta BEL, Filho JBG, et al (2019) Use of anaerobic co-digestion as an alternative to add value to sugarcane biorefinery wastes. Bioresour Technol 287:121443. https://doi.org/10.1016/j.biortech.2019.121443

7. Djalma Nunes Ferraz Júnior A, Koyama MH, de Araújo Júnior MM, Zaiat M (2016) Thermophilic anaerobic digestion of raw sugarcane vinasse. Renew Energy 89:245–252. https://doi.org/10.1016/j.renene.2015.11.064

8. Moraes BS, Zaiat M, Bonomi A (2015) Anaerobic digestion of vinasse from sugarcane ethanol production in Brazil: Challenges and perspectives. Renew Sustain Energy Rev 44:888–903. https://doi.org/10.1016/j.rser.2015.01.023

9. Fuess LT, Kiyuna LSM, Ferraz ADN, et al (2017) Thermophilic two-phase anaerobic digestion using an innovative fixed-bed reactor for enhanced organic matter removal and bioenergy recovery from sugarcane vinasse. Appl Energy 189:480–491. https://doi.org/10.1016/j.apenergy.2016.12.071

10. Menandro LMS, Cantarella H, Franco HCJ, et al (2017) Comprehensive assessment of sugarcane straw: implications for biomass and bioenergy production. Biofuels, Bioprod Biorefining 11:488–504. https://doi.org/10.1002/bbb.1760

11. Khaire KC, Moholkar VS, Goyal A (2021) Bioconversion of sugarcane tops to bioethanol and other value added products: An overview. Mater Sci Energy Technol 4:54–68. https://doi.org/10.1016/j.mset.2020.12.004

12. Brenelli LB, Figueiredo FL, Damasio A, et al (2020) An integrated approach to obtain xylo-oligosaccharides from sugarcane straw: from lab to pilot scale. Bioresour Technol 123637. https://doi.org/10.1016/j.biortech.2020.123637

13. Janke L, Leite A, Batista K, et al (2016) Optimization of hydrolysis and volatile fatty acids production from sugarcane filter cake: Effects of urea supplementation and sodium hydroxide pretreatment. Bioresour Technol 199:235–244. https://doi.org/10.1016/j.biortech.2015.07.117

14. Wongarmat W, Reungsang A, Sittijunda S, Chu CY (2021) Anaerobic codigestion of biogas effluent and sugarcane filter cake for methane production. Biomass Convers Biorefinery. https://doi.org/10.1007/s13399-021-01305-3

15. Tellechea FRF, Martins MA, da Silva AA, et al (2016) Use of sugarcane filter cake and nitrogen, phosphorus and potassium fertilization in the process of bioremediation of soil contaminated with diesel. Environ Sci Pollut Res 23:18027–18033. https://doi.org/10.1007/s11356-016-6965-x

16. Hernández-Pérez AF, de Arruda PV, Felipe M das G de A (2016) Sugarcane straw as a feedstock for xylitol production by *Candida guilliermondii* FTI 20037. Brazilian J Microbiol 47:489–496. https://doi.org/10.1016/j.bjm.2016.01.019

17. Hagos K, Zong J, Li D, et al (2017) Anaerobic co-digestion process for biogas production: Progress, challenges and perspectives. Renew. Sustain. Energy Rev.

18. Janke L, Leite AF, Nikolausz M, et al (2016) Comparison of start-up strategies and process performance during semi-continuous anaerobic digestion of sugarcane filter cake co-digested with bagasse. Waste Manag 48:199–208. https://doi.org/10.1016/j.wasman.2015.11.007

19. Raposo F, Fernández-Cegrí V, de la Rubia MA, et al (2011) Biochemical methane potential (BMP) of solid organic substrates: Evaluation of anaerobic biodegradability using data from an international interlaboratory study. J Chem Technol Biotechnol 86:1088–1098. https://doi.org/10.1002/jctb.2622

20. Hobbs SR, Landis AE, Rittmann BE, et al (2018) Enhancing anaerobic digestion of food waste through biochemical methane potential assays at different substrate: inoculum ratios. Waste Manag 71:612–617. https://doi.org/10.1016/j.wasman.2017.06.029

21. Sluiter A, Hames B, Ruiz R, et al (2008) Determination of sugars, byproducts, and degradation products in liquid fraction process samples, Technical Report NREL/TP-510-42623

22. Rodriguez-Chiang L, Llorca J, Dahl O (2016) Anaerobic co-digestion of acetate-rich with lignin-rich wastewater and the effect of hydrotalcite addition. Bioresour Technol 218:84–91. https://doi.org/10.1016/j.biortech.2016.06.074

23. Triolo JM, Pedersen L, Qu H, Sommer SG (2012) Biochemical methane potential and anaerobic biodegradability of non-herbaceous and herbaceous phytomass in biogas production. Bioresour Technol 125:226–232. https://doi.org/10.1016/j.biortech.2012.08.079

24. VDI 4630 (2006) Fermentation of organic materials. Characterization of the substrate, sampling, collection of material data, fermentation tests.

25. Boušková A, Dohányos M, Schmidt JE, Angelidaki I (2005) Strategies for changing temperature from mesophilic to thermophilic conditions in anaerobic CSTR reactors treating sewage sludge. Water Res 39:1481–1488. https://doi.org/10.1016/j.watres.2004.12.042

26. Filer J, Ding HH, Chang S (2019) Biochemical Methane Potential (BMP) Assay Method for Anaerobic Digestion Research. Water 11:29. https://doi.org/10.3390/w11050921

27. Mockaitis G, Bruant G, Guiot SR, et al (2020) Acidic and thermal pre-treatments for anaerobic digestion inoculum to improve hydrogen and volatile fatty acid production using xylose as the substrate. Renew Energy 145:1388–1398. https://doi.org/10.1016/j.renene.2019.06.134

28. APHA, AWWA W (2012) Standard Methods for the Examination of Water and Wastewater, twenty-sec. Washington, DC.

29. Sluiter JB, Chum H, Gomes AC, et al (2016) Evaluation of Brazilian sugarcane bagasse characterization: An interlaboratory comparison study. J AOAC Int 99:579–585. https://doi.org/10.5740/jaoacint.15-0063

30. Janke L, Weinrich S, Leite AF, et al (2017) Optimization of semi-continuous anaerobic digestion of sugarcane straw co-digested with filter cake: Effects of macronutrients supplementation on conversion kinetics. Bioresour Technol 245:35–43. https://doi.org/10.1016/j.biortech.2017.08.084

31. Friehe J, Weiland P, Schattauer (2010) Fundamentals of anaerobic digestion In: FNR (Fachagentur Nachwachsende Rohstoffe e.V.) (Ed) Guide to Biogas—From Production to Use. FNR: Germany, pp 21–31.

32. Tan L, Zhong J, Jin YL, et al (2020) Production of bioethanol from unwashed-pretreated rapeseed straw at high solid loading. Bioresour Technol 303:122949. https://doi.org/10.1016/j.biortech.2020.122949

33. Junqueira TL, Moraes BS, Gouveia VLR, et al (2015) Use of VSB to Plan Research Programs and Public Policies. In: Bonomi A, Cavalett O, Pereira da Cunha M, Lima M (eds) Virtual Biorefinery: An Optimization Strategy for Renewable Carbon Valorization. Springer, Campinas, pp 257–282

34. Pérez-Pimienta JA, Icaza-Herrera JPA, Méndez-Acosta HO, et al (2020) Bioderived ionic liquid-based pretreatment enhances methane production from: Agave tequilana bagasse. RSC Adv 10:14025–14032. https://doi.org/10.1039/d0ra01849j

35. Kamdem I, Hiligsmann S, Vanderghem C, et al (2018) Enhanced Biogas Production During Anaerobic Digestion of Steam-Pretreated Lignocellulosic Biomass from Williams Cavendish Banana Plants. Waste and Biomass Valorization 9:175–185. https://doi.org/10.1007/s12649-016-9788-6

36. Demirel B, Scherer P (2008) The roles of acetotrophic and hydrogenotrophic methanogens during anaerobic conversion of biomass to methane: A review. Rev Environ Sci Biotechnol 7:173–190. https://doi.org/10.1007/s11157-008-9131-1

37. Wang Y, Zhang Y, Wang J, Meng L (2009) Effects of volatile fatty acid concentrations on methane yield and methanogenic bacteria. Biomass and Bioenergy. https://doi.org/10.1016/j.biombioe.2009.01.007

38. Menon A, Wang JY, Giannis A (2017) Optimization of micronutrient supplement for enhancing biogas production from food waste in two-phase thermophilic anaerobic digestion. Waste Manag 59:465–475. https://doi.org/10.1016/j.wasman.2016.10.017

39. Ahn J-H, Do TH, Kim SD, Hwang S (2006) The effect of calcium on the anaerobic digestion treating swine wastewater. Biochem Eng J 30:33–38. https://doi.org/https://doi.org/10.1016/j.bej.2006.01.014

40. Romero-Güiza MS, Mata-Alvarez J, Chimenos JM, Astals S (2016) The effect of magnesium as activator and inhibitor of anaerobic digestion. Waste Manag 56:137–142. https://doi.org/https://doi.org/10.1016/j.wasman.2016.06.037

41. Koster IW, Rinzema A, de Vegt AL, Lettinga G (1986) Sulfide inhibition of the methanogenic activity of granular sludge at various pH-levels. Water Res 20:1561–1567. https://doi.org/10.1016/0043-1354(86)90121-1

42. Chen Y, Cheng JJ, Creamer KS (2008) Inhibition of anaerobic digestion process: a review. Bioresour Technol 99:4044–64. https://doi.org/10.1016/j.biortech.2007.01.057

43. Issah A, Kabera T, Kemausuor F (2020) Biomass and Bioenerg Biogas optimisation processes and effluent quality □: A review. Biomass and Bioenergy 133:105449. https://doi.org/10.1016/j.biombioe.2019.105449

44. Bhattacharya SK, Uberoi V, Madura RL, Haghighi-Podeh MR (1995) Effect of Cobalt on Methanogenesis. Environ Technol 16:271–278. https://doi.org/10.1080/09593331608616269

45. Zieliński M, Kisielewska M, Dębowski M, Elbruda K (2019) Effects of Nutrients Supplementation on Enhanced Biogas Production from Maize Silage and Cattle Slurry Mixture. Water Air Soil Pollut 230:. https://doi.org/10.1007/s11270-019-4162-5

46. Zhang J, Zou W, Li Y, et al (2015) Plant Science Silica distinctively affects cell wall features and lignocellulosic saccharification with large enhancement on biomass production in. Plant Sci 239:84–91. https://doi.org/10.1016/j.plantsci.2015.07.014

47. Rasi S, Seppälä M, Rintala J (2013) Organic silicon compounds in biogases produced from grass silage, grass and maize in laboratory batch assays. Energy 52:137–142. https://doi.org/10.1016/j.energy.2013.01.015

48. Feijoo G, Soto M, Méndez R, Lema JM (1995) Sodium inhibition in the anaerobic digestion process: Antagonism and adaptation phenomena. Enzyme Microb Technol 17:180–188. https://doi.org/10.1016/0141-0229(94)00011-F

49. Qiang H, Niu Q, Chi Y, Li Y (2013) Trace metals requirements for continuous thermophilic methane fermentation of high-solid food waste. Chem Eng J 222:330–336. https://doi.org/10.1016/j.cej.2013.02.076

50. Holliger C, Alves M, Andrade D, et al (2016) Towards a standardization of biomethane potential tests. Water Sci Technol 74:2515–2522. https://doi.org/10.2166/wst.2016.336

51. Galaction AI, Cascaval D, Oniscu C, Turnea M (2004) Prediction of oxygen mass transfer coefficients in stirred bioreactors for bacteria, yeasts and fungus broths. Biochem Eng J 20:85–94. https://doi.org/10.1016/j.bej.2004.02.005

52. Rabelo SC, Andrade RR, Maciel Filho R, Costa AC (2014) Alkaline hydrogen peroxide pretreatment, enzymatic hydrolysis and fermentation of sugarcane bagasse to ethanol. Fuel 136:349–357. https://doi.org/10.1016/j.fuel.2014.07.033

53. Tramontina R, Brenelli LB, Sousa A, et al (2020) Designing a cocktail containing redox enzymes to improve hemicellulosic hydrolysate fermentability by microorganisms. Enzyme Microb Technol 135:109490. https://doi.org/10.1016/j.enzmictec.2019.109490

54. Khan MU, Ahring BK (2019) Lignin degradation under anaerobic digestion: Influence of lignin modifications -A review. Biomass and Bioenerg 128:. https://doi.org/10.1016/j.biombioe.2019.105325

55. Mulat DG, Horn SJ (2018) Chapter 14: Biogas Production from Lignin via Anaerobic Digestion. RSC Energy Environ Ser 2018-Janua:391–412. https://doi.org/10.1039/9781788010351-00391

56. Gonzalez-Gil G, Seghezzo L, Lettinga G, et al (2001) Kinetics and mass-transfer phenomena in anaerobic granular sludge. Biotechnol Bioeng 73:125–134. https://doi.org/10.1002/bit.1044

57. Krishania M, Vijay VK, Chandra R (2013) Methane fermentation and kinetics of wheat straw pretreated substrates co-digested with cattle manure in batch assay. Energy 57:359–367. https://doi.org/10.1016/j.energy.2013.05.028

58. Ganesh Kumar A, Sekaran G, Krishnamoorthy S (2006) Solid state fermentation of Achras zapota lignocellulose by Phanerochaete chrysosporium. Bioresour Technol 97:1521–1528. https://doi.org/10.1016/j.biortech.2005.06.015

59. Barakat A, Monlau F, Steyer JP, Carrere H (2012) Effect of lignin-derived and furan compounds found in lignocellulosic hydrolysates on biomethane production. Bioresour Technol 104:90–99. https://doi.org/10.1016/j.biortech.2011.10.060

60. Kim JY, Park J, Hwang H, et al (2015) Catalytic depolymerization of lignin macromolecule to alkylated phenols over various metal catalysts in supercritical tert-butanol. J Anal Appl Pyrolysis 113:99–106. https://doi.org/10.1016/j.jaap.2014.11.011

61. Demirel B, Yenigün O (2002) Two-phase anaerobic digestion processes: A review. J Chem Technol Biotechnol 77:743–755. https://doi.org/10.1002/jctb.630

62. Gu Y, Chen X, Liu Z, et al (2014) Effect of inoculum sources on the anaerobic digestion of rice straw. Bioresour Technol 158:149–155. https://doi.org/10.1016/j.biortech.2014.02.011

63. Quintero M, Castro L, Ortiz C, et al (2012) Enhancement of starting up anaerobic digestion of lignocellulosic substrate: Fique’s bagasse as an example. Bioresour Technol 108:8–13. https://doi.org/10.1016/j.biortech.2011.12.052

