## Supplementary material for "Use of lignocellulosic residue from second-generation ethanol production to enhance methane production through co-digestion": Original dataset paper

### Dataset of anaerobic digestion and co-digestion of residues from 1G2G sugarcane: Biochemical Methane Potential data and kinetics fitting

Maria Paula C. Volpi*[a,b], Brenno V. M. Lima,[b,c], Gustavo Mockaitis[b], Bruna S. Moraes[a,b]

[a]Interdisciplinary Center of Energy Planning, University of Campinas (NIPE/UNICAMP), R, Cora Coralina, 330 - Cidade Universitária, Campinas - SP, 13083-896, Brazil.

[b]Interdisciplinary Research Group on Biotechnology Applied to the Agriculture and the Environment (GBMA), School of Agricultural Engineering (FEAGRI), University of Campinas (UNICAMP), Av, Candido Rondon, 501 - Cidade Universitária, Campinas - SP, 13083-875, Brazil.

[c]School of Mechanical Engineering (FEM), University of Campinas (UNICAMP), Rua Mendeleyev, 200 - Cidade Universitária, Campinas – SP, 13083-860, Brazil.

*Corresponding author: mcardealvolpi@gmail,com; phone (+55) 19 98367-2506

**Abstract**

This paper presents the raw data of the Biochemical Methane Potential (BMP) essay for vinasse, filter cake (residues from 1G ethanol production), and deacetylation liquor (residue from 2G ethanol production). Those essays evaluated the BMP of each sole residue and their co-digestion. Experiments evaluated two different inocula (from a sugarcane vinasse treatment plant and a poultry slaughterhouse) to obtain each experiment's BMP. Data presented in this article is the biogas production, volatile solids amount, and $CH_4$ concentration for each studied condition. Kinetic modeling for methane production was also included in this dataset. All data (spreadsheets and modeling files) are available at Mendeley Data (DOI: 10.17632/6bm43w2dvt.1).

**Keywords: Biochemical Methane Potential, Co-digestion, Sugarcane Biorefinery, 2G ethanol residue**

**Specification Tables**

| | |
|---|---|
| **Subject** | Biotechnology |
| **Specific subject area** | Bioenergy and Biofuels |
| **Type of data** | Tables<br>Graphs<br>Spreadsheets |
| **How data were acquired** | Experiments were conducted in single batches using 250 mL Duran® flasks, containing 40% headspace, The volume of biogas was measured daily using Gastight Hamilton Super Syringe (1L), and 3 mL of gas was removed for analysis in Gas chromatography using the |

| | |
|---|---|
| | Construmaq MOD, U-13, São Carlos (GC), For data analysis and treatment was used computer, |
| **Data format** | Raw data, kinetic model fitting |
| **Parameters for data collection** | The collected data were biogas production (mL), methane molar/volumetric fraction (% $CH_4$), and volatile solids (mg $L^{-1}$), The biogas was collected at room temperature, as well as the analysis of volatile solids, The reading of the concentration of $CH_4$ in the GC was made in 22°C, |
| **Description of data collection** | Data were collected from 250 mL flasks that had a butyl cap for sealing, A syringe was placed in each bottle to remove the biogas from the headspace |
| **Data source location** | Environment and Sanitation Laboratory, School of Agricultural Engineering, University of Campinas (LMAS/FEAGRI/UNICAMP) Candido Rondon Avenue 501, Cidade Universitária, 13,083-875, Campinas, SP – Brazil (22º49’12,6”S – 47º03’39,5”W), |

| | |
|---|---|
| **Data accessibility** | All spreadsheets and kinetic data are available at Mendeley Data, through DOI:10,17632/6bm43w2dvt,1<br><br>Related research Article: |

**Value of the Data**

- Biochemical Methane Potential of residues is important to know how much CH4 the residues from the 1G2G sugarcane mill can produce,
- This data can be used by researchers in anaerobic digestion areas or co-digestion and by sugarcane mills,
- All data presented in this paper can be used for comparison with other experiments with anaerobic co-digestion

#### 1. Data Description

Biochemical methane potential (BMP) is a technique to determine the production of $CH_4$ of an organic substrate during its anaerobic decomposition. This test is a straightforward and reliable method to obtain the conversion rate of organic matter to $CH_4$ [1] and determine the maximum capacity of $CH_4$ production. BMP technique is used to design biodigesters and bioreactors for biogas production and waste management, especially when working with substrates not yet subjected to anaerobic digestion. BMP provides information that could predict the behavior of substrate degradation and its potential to produce $CH_4$.

The present dataset comprises raw data from BMP tests such as biogas production, methane contents, and volatile solids (VS). BMP data was acquired from an entire experimental setup designed to evaluate the effect of two different inoculants and different combinations of residues from a sugarcane mill producing both 1G and 2G ethanol. Experiments used the filter cake, vinasse, and deacetylation residue as substrates.

Accumulated biogas production of each experiment used to calculate the BMP are described in Table 2 and Table 3 (regarding experiment A and experiment B, respectively). Table 4 shows the average $CH_4$ molar fraction for both experiments A and B. Tables 5 and 6 show the accumulated $CH_4$ production values for experiments A and B, respectively). Tables 7 and 8 show the kinetic model's values to produce $CH_4$ in each of the tests, in respect of time.

All experiments were carried out in triplicate. The data depicted in this data article are the average values of these triplicates. All the raw data for each replica and the calculations performed are already published and available in the Mendeley Data (DOI: 10,17632/6bm43w2dvt,1).

#### 2. Experimental Design, Materials, and Methods

##### 2.1 Experimental arrangement

The experimental design used to evaluate the BMP of sole residues and their combination in co-digestion is depicted in Table 1.

Table 1 – Experimental Biochemical Methane Potential (BMP) design for this study.

| Assay | Bulk Composition (weight-based %, in terms of VS) | | | | |
|---|---|---|---|---|---|
| | Experiment A – Inoculum from vinasse processing digestor | | | | |
| | Vinasse | Filter Cake | Liquor | Cellulose | Inoculum |
| A1 | 33 | - | - | - | 66 |
| A2 | - | 33 | - | - | 66 |

| | | | | | |
|---|---|---|---|---|---|
| A3 | - | - | 33 | - | 66 |
| A4* | 11 | 11 | 11 | - | 66 |
| A+ | - | - | - | 33 | 66 |
| A- | - | - | - | - | 100 |
| Assay | Experiment B – Inoculum from poultry slaughterhouse wastewater digestor | | | | |
| B1 | 33 | - | - | - | 66 |
| B2 | - | 33 | - | - | 66 |
| B3 | - | - | 33 | - | 66 |
| B4 | 11 | 11 | 11 | - | 66 |
| B5* | - | - | - | 33 | 66 |
| B6* | - | - | - | - | 100 |
| B7* | - | 17 | 17 | - | 66 |
| B+ | - | - | - | 33 | 66 |
| B- | - | - | - | - | 100 |

*co-digestion assays, + positive control (cellulose as substrate); - negative control (without substrate).

The substrates used were vinasse, filter cake (both from 1G ethanol production) obtained from Iracema Mill (São Martinho group), Iracemápolis municipality, São Paulo state, Brazil (22°35’17.6”S 47°31’51.5” W). Deacetylation liquor was obtained from a sugarcane straw pretreatment (2G ethanol), that was produced in the National Laboratory of Biorenewables (LNBR/CNPEM). Experiment A used an inoculum from Iracema mill (São Martinho group), obtained from the sludge of a mesophilic reactor (BIOPAQ®ICX - Paques), treating sugarcane vinasse. Experiment 2 was inoculated with a sludge obtained from an Upflow Anaerobic Sludge Blanket UASB bioreactor treating wastewater from a poultry slaughterhouse (Ideal slaughterhouse company), at Pereiras municipality, São Paulo state, Brazil (23°05’10.5”S 47°58’58.9” W).

According to the Standard Methods for the Examination of Water and Wastewater [2], organic matter content in the substrates was determined using the 5220B method for Chemical Oxygen Demand (COD) determination and 2540 method for the solid series.

The solids series methodology considered the concentration of total (TS, mg $L^{-1}$), and volatile (VS, mg $L^{-1}$) solids.

All experiments were conducted in triplicates of single batches using 250 mL Duran® flasks as bioreactors closed with a pierceable isobutylene isoprene rubber septum, and stored in an Ethickt echnology (411-FPD) incubator at thermophilic condition (55°C), followed the VDI 4630 [3] protocol and Triolo et al, [1]. Two BMP assay were performed, The first assay (experiment A) used the mill inoculum, and flasks were assembled for each residue individually (vinasse, filter cake, and deacetylation liquor), a flask co-digestion of the 3 residues together, positive control using cellulose and negative control with a flask of inoculum only. In experiment B, poultry slaughterhouse inoculum was used, the same flasks were made as in experiment A, but additionally flasks with co-digestion of the residues two by two, Table 1 best exemplifies the experimental design.

The experiment was 2:1 inoculum to substrate ratio (w/w, in terms of Volatile Solids-VS) added to each flask, thus ensuring excess of inoculum to consume all the organic matter of the substrate and achieving its maximum experimental $CH_4$ production, The pH of solution flasks was corrected to neutrality by adding solutions of NaOH (0,5 M) or $H_2SO_4$ (1 M) when necessary, Nitrogen ($N_2$) gas was fluxed into the liquid medium for 10 min and into the headspace for 5 min after closing the flasks, The headspace was kept at 40%, Biogas was collected from the headspace over the days by using a Gastight Hamilton Super Syringe (1 L) through the flasks' rubber septum, and was read $CH_4$ concentration in Gas chromatography (Construmaq MOD, U-13, São Carlos),

Table 2, Average biogas accumulation for experiment 1

| Time (days) | Vinasse (mL biogas) | Deacetylation Liquor (mL biogas) | Filter Cake (mL biogas) | Co-digestion (mL biogas) | Cellulose (mL biogas) |
|---|---|---|---|---|---|
| 1 | 72 | 50 | 70 | 83 | 69 |
| 2 | 133 | 91 | 131 | 145 | 121 |
| 3 | 191 | 161 | 185 | 200 | 173 |
| 4 | 245 | 173 | 240 | 255 | 225 |
| 5 | 299 | 195 | 297 | 307 | 290 |
| 6 | 356 | 228 | 348 | 360 | 352 |
| 7 | 408 | 251 | 406 | 413 | 411 |
| 8 | 461 | 275 | 457 | 465 | 465 |
| 9 | 513 | 301 | 508 | 517 | 518 |
| 10 | 559 | 341 | 561 | 568 | 572 |
| 12 | 611 | 378 | 611 | 621 | 627 |
| 14 | 663 | 420 | 665 | 674 | 683 |
| 16 | 714 | 433 | 720 | 727 | 740 |
| 18 | 770 | 446 | 792 | 779 | 824 |
| 20 | 822 | 490 | 848 | 831 | 881 |
| 23 | 872 | 540 | 924 | 890 | 938 |
| 26 | 925 | 557 | 995 | 958 | 999 |
| 29 | 975 | 576 | 1056 | 1017 | 1055 |
| 30 | 1007 | 630 | 1133 | 1102 | 1117 |
| 33 | 1057 | 690 | 1217 | 1196 | 1188 |
| 35 | 1101 | 857 | 1291 | 1272 | 1248 |
| 38 | 1162 | 911 | 1359 | 1369 | 1314 |
| 41 | 1230 | 970 | 1423 | 1440 | 1379 |
| 43 | 1301 | 1027 | 1490 | 1509 | 1443 |
| 46 | 1352 | 1079 | 1530 | 1572 | 1508 |
| 49 | 1394 | 1137 | 1597 | 1636 | 1575 |
| 51 | 1446 | 1205 | 1658 | 1696 | 1631 |
| 55 | 1495 | 1269 | 1715 | 1753 | 1685 |
| 58 | 1543 | 1340 | 1770 | 1809 | 1739 |
| 60 | 1593 | 1400 | 1832 | 1867 | 1800 |
| 61 | 1644 | 1453 | 1885 | 1918 | 1850 |
| 63 | 1702 | 1517 | 1948 | 1977 | 1910 |
| 65 | 1757 | 1579 | 2005 | 2027 | 1960 |
| 67 | 1807 | 1639 | 2065 | 2086 | 2017 |
| 68 | 1860 | 1701 | 2127 | 2146 | 2074 |
| 70 | 1900 | 1745 | 2167 | 2180 | 2114 |
| 73 | 1958 | 1782 | 2232 | 2197 | 2161 |
| 76 | 2026 | 1837 | 2292 | 2232 | 2205 |
| 79 | 2095 | 1892 | 2356 | 2249 | 2251 |
| 80 | 2167 | 1947 | 2419 | 2299 | 2295 |

Table 3, Average biogas accumulation for experiment 2

| Time (days) | Vinasse (mL biogas) | Deacetylation Liquor (mL biogas) | Filter Cake (mL biogas) | Co-digestion (mL biogas) | Cellulose (mL biogas) | Filter Cake + vinasse (mL biogas) | Filter Cake + Deacetylation Liquor (mL biogas) | Vinasse + Deacetylation Liquor |
|---|---|---|---|---|---|---|---|---|
| 1 | 75 | 63 | 57 | 75 | 51 | 73 | 65 | 71 |
| 2 | 148 | 125 | 108 | 143 | 92 | 141 | 121 | 140 |
| 3 | 214 | 186 | 148 | 200 | 135 | 200 | 173 | 203 |
| 4 | 270 | 239 | 190 | 250 | 174 | 251 | 223 | 255 |
| 5 | 323 | 287 | 236 | 300 | 218 | 301 | 273 | 315 |
| 6 | 383 | 346 | 292 | 358 | 282 | 361 | 340 | 371 |
| 7 | 433 | 376 | 344 | 403 | 347 | 412 | 396 | 420 |
| 8 | 487 | 422 | 399 | 452 | 405 | 461 | 454 | 463 |
| 9 | 535 | 474 | 459 | 499 | 458 | 511 | 515 | 512 |
| 10 | 567 | 534 | 539 | 551 | 536 | 567 | 653 | 555 |
| 12 | 612 | 597 | 600 | 602 | 585 | 618 | 720 | 600 |
| 14 | 660 | 719 | 684 | 660 | 626 | 668 | 806 | 650 |
| 16 | 709 | 762 | 767 | 748 | 666 | 728 | 892 | 697 |
| 18 | 756 | 786 | 867 | 827 | 706 | 798 | 971 | 728 |
| 20 | 803 | 928 | 955 | 919 | 748 | 918 | 1035 | 777 |
| 23 | 848 | 967 | 1024 | 960 | 788 | 959 | 1109 | 829 |
| 26 | 894 | 1020 | 1077 | 999 | 832 | 1065 | 1176 | 882 |
| 29 | 943 | 1072 | 1147 | 1041 | 875 | 1157 | 1232 | 939 |
| 30 | 967 | 1114 | 1185 | 1091 | 903 | 1207 | 1278 | 996 |
| 33 | 1007 | 1165 | 1250 | 1133 | 940 | 1277 | 1334 | 1091 |
| 35 | 1020 | 1220 | 1314 | 1190 | 980 | 1357 | 1393 | 1174 |
| 38 | 1061 | 1268 | 1375 | 1247 | 1032 | 1418 | 1448 | 1244 |
| 41 | 1103 | 1301 | 1417 | 1295 | 1073 | 1477 | 1495 | 1305 |
| 43 | 1144 | 1346 | 1460 | 1341 | 1115 | 1525 | 1544 | 1356 |

| | | | | | | | | |
|---|---|---|---|---|---|---|---|---|
| 46 | 1190 | 1392 | 1505 | 1389 | 1169 | 1583 | 1593 | 1409 |
| 49 | 1200 | 1432 | 1535 | 1437 | 1239 | 1634 | 1643 | 1459 |
| 51 | 1238 | 1480 | 1577 | 1487 | 1314 | 1685 | 1690 | 1507 |
| 55 | 1283 | 1509 | 1617 | 1534 | 1392 | 1735 | 1733 | 1557 |
| 58 | 1329 | 1551 | 1665 | 1584 | 1496 | 1780 | 1782 | 1589 |
| 60 | 1359 | 1597 | 1703 | 1628 | 1592 | 1831 | 1830 | 1635 |
| 61 | 1409 | 1612 | 1748 | 1677 | 1699 | 1879 | 1880 | 1685 |
| 63 | 1460 | 1657 | 1796 | 1727 | 1856 | 1929 | 1935 | 1736 |
| 65 | 1512 | 1707 | 1852 | 1778 | 1938 | 1982 | 1983 | 1786 |
| 67 | 1551 | 1707 | 1905 | 1778 | 2022 | 2020 | 2001 | 1822 |
| 68 | 1602 | 1759 | 1964 | 1834 | 2168 | 2062 | 2042 | 1840 |
| 70 | 1670 | 1818 | 2023 | 1892 | 2308 | 2121 | 2100 | 1898 |
| 73 | 1745 | 1876 | 2091 | 1952 | 2408 | 2186 | 2161 | 1959 |
| 76 | 1823 | 1894 | 2148 | 2012 | 2502 | 2240 | 2219 | 2019 |
| 79 | 1919 | 1922 | 2208 | 2050 | 2616 | 2296 | 2279 | 2037 |
| 80 | 1997 | 1939 | 2262 | 2106 | 2692 | 2356 | 2335 | 2095 |
| 83 | 2107 | 1948 | 2326 | 2165 | 2801 | 2416 | 2389 | 2153 |
| 86 | 2266 | 1983 | 2384 | 2279 | 2882 | 2474 | 2446 | 2211 |
| 89 | 2268 | 1990 | 2453 | 2317 | 2957 | 2534 | 2501 | 2268 |
| 92 | 2272 | 1990 | 2507 | 2367 | 3046 | 2608 | 2554 | 2325 |
| 95 | 2274 | 1995 | 2563 | 2590 | 3114 | 2663 | 2614 | 2379 |
| 99 | 2277 | 1995 | 2617 | 2685 | 3169 | 2723 | 2674 | 2433 |
| 102 | 2279 | 1995 | 2673 | 2799 | 3226 | 2781 | 2731 | 2486 |
| 104 | 2281 | 1995 | 2730 | 2875 | 3281 | 2840 | 2785 | 2542 |
| 107 | 2281 | 2012 | 2786 | 2930 | 3341 | 2897 | 2839 | 2599 |
| 109 | 2282 | 2016 | 2843 | 2994 | 3405 | 2961 | 2891 | 2656 |
| 112 | 2286 | 2034 | 2900 | 3044 | 3513 | 3021 | 2929 | 2713 |
| 113 | 2287 | 2034 | 2957 | 3077 | 3570 | 3082 | 2959 | 2770 |
| 114 | 2288 | 2034 | 3014 | 3111 | 3627 | 3143 | 2989 | 2827 |
| 115 | 2288 | 2035 | 3033 | 3127 | 3684 | 3203 | 3019 | 2881 |

Table 4, Average of $CH_4$ concentration in both experiments

| Assay | Average of $CH_4$ concentration (%) |
|---|---|
| Experiment 1 | |
| Vinasse | 57.48 |
| Filter Cake | 58.14 |
| Deacetylation Liquor | 67.50 |
| Co-digestion | 64.67 |
| Cellulose | 69.91 |
| Experiment 2 | |
| Vinasse | 57.50 |
| Filter Cake | 58.20 |
| Deacetylation Liquor | 67.55 |
| Co-digestion | 64.70 |
| Cellulose | 69.95 |
| Vinasse + filter cake | 76.37 |
| Vinasse + Deacetylation Liquor | 70.86 |
| Deacetylation liquor + filter cake | 61.58 |

Table 5, Average production of $CH_4$ from the BMP assays in experiment 1

| Time (days) | Vinasse (NmLCH$_4$ gVS$^{-1}$) | Deacetylation Liquor (NmLCH$_4$ gVS$^{-1}$) | Filter Cake (NmLCH$_4$ gVS$^{-1}$) | Co-digestion (NmLCH$_4$ gVS$^{-1}$) | Cellulose (NmLCH$_4$ gVS$^{-1}$) |
|---|---|---|---|---|---|
| 1 | 17 | 2 | 3 | 30 | 2 |
| 2 | 33 | 2 | 11 | 50 | 2 |
| 3 | 49 | 48 | 13 | 63 | 3 |
| 4 | 61 | 13 | 18 | 77 | 5 |
| 5 | 72 | 13 | 23 | 87 | 19 |
| 6 | 87 | 13 | 24 | 98 | 30 |
| 7 | 97 | 13 | 31 | 109 | 39 |
| 8 | 108 | 13 | 31 | 120 | 44 |
| 9 | 118 | 13 | 32 | 130 | 46 |
| 10 | 121 | 13 | 35 | 139 | 50 |
| 12 | 132 | 13 | 35 | 151 | 56 |
| 14 | 142 | 13 | 39 | 162 | 61 |

| | | | | | |
|---|---|---|---|---|---|
| 16 | 152 | 13 | 44 | 174 | 69 |
| 18 | 162 | 13 | 60 | 181 | 100 |
| 20 | 171 | 13 | 65 | 192 | 107 |
| 23 | 180 | 13 | 87 | 210 | 114 |
| 26 | 190 | 13 | 105 | 239 | 125 |
| 29 | 200 | 13 | 116 | 259 | 132 |
| 30 | 203 | 13 | 153 | 324 | 161 |
| 33 | 212 | 13 | 184 | 386 | 184 |
| 35 | 210 | 94 | 201 | 421 | 190 |
| 38 | 218 | 102 | 206 | 470 | 193 |
| 41 | 225 | 109 | 198 | 481 | 185 |
| 43 | 232 | 112 | 191 | 483 | 172 |
| 46 | 240 | 138 | 180 | 505 | 184 |
| 49 | 233 | 165 | 189 | 522 | 196 |
| 51 | 241 | 194 | 196 | 540 | 198 |
| 55 | 247 | 224 | 201 | 555 | 200 |
| 58 | 252 | 257 | 204 | 568 | 204 |
| 60 | 254 | 287 | 210 | 581 | 207 |
| 61 | 264 | 308 | 212 | 589 | 206 |
| 63 | 274 | 339 | 218 | 601 | 210 |
| 65 | 285 | 369 | 220 | 607 | 207 |
| 67 | 293 | 402 | 228 | 626 | 214 |
| 68 | 304 | 469 | 237 | 646 | 221 |
| 70 | 325 | 487 | 255 | 666 | 245 |
| 73 | 351 | 491 | 279 | 651 | 259 |
| 76 | 383 | 525 | 296 | 659 | 270 |
| 79 | 428 | 569 | 326 | 639 | 278 |
| 80 | 487 | 605 | 363 | 660 | 282 |

Table 6, Average production of $CH_4$ from the BMP assays in experiment 2

| Time (days) | Vinasse (NmLCH$_4$ gVS$^{-1}$) | Deacetylation Liquor (NmLCH$_4$ gVS$^{-1}$) | Filter Cake (NmLCH$_4$ gVS$^{-1}$) | Co-digestion (NmLCH$_4$ gVS$^{-1}$) | Cellulose (NmLCH$_4$ gVS$^{-1}$) | Filter Cake + vinasse (NmLCH$_4$ gVS$^{-1}$) | Filter Cake + Deacetylation Liquor (NmLCH$_4$ gVS$^{-1}$) | Vinasse + Deacetylation Liquor (NmLCH$_4$ gVS$^{-1}$) |
|---:|---:|---:|---:|---:|---:|---:|---:|---:|
| 1 | 17 | 27 | 2 | 13 | 0 | 12 | 15 | 22 |
| 2 | 33 | 53 | 4 | 25 | 1 | 23 | 29 | 44 |
| 3 | 49 | 80 | 5 | 34 | 1 | 33 | 41 | 64 |
| 4 | 61 | 102 | 6 | 41 | 2 | 40 | 52 | 80 |
| 5 | 72 | 121 | 7 | 49 | 2 | 47 | 63 | 99 |
| 6 | 87 | 147 | 12 | 60 | 8 | 58 | 83 | 117 |
| 7 | 97 | 156 | 14 | 67 | 13 | 65 | 96 | 131 |
| 8 | 108 | 177 | 18 | 74 | 17 | 73 | 112 | 144 |
| 9 | 118 | 198 | 22 | 82 | 20 | 80 | 128 | 158 |
| 10 | 121 | 224 | 29 | 90 | 27 | 89 | 173 | 170 |
| 12 | 132 | 253 | 33 | 99 | 29 | 97 | 193 | 183 |
| 14 | 142 | 316 | 41 | 109 | 30 | 105 | 218 | 198 |
| 16 | 152 | 332 | 51 | 127 | 31 | 115 | 244 | 213 |
| 18 | 162 | 336 | 65 | 143 | 32 | 128 | 267 | 219 |
| 20 | 171 | 413 | 74 | 161 | 32 | 154 | 284 | 234 |
| 23 | 180 | 426 | 79 | 165 | 33 | 157 | 304 | 249 |
| 26 | 190 | 449 | 77 | 170 | 35 | 182 | 323 | 266 |
| 29 | 200 | 471 | 81 | 175 | 36 | 200 | 338 | 284 |
| 30 | 203 | 492 | 84 | 184 | 36 | 210 | 351 | 304 |
| 33 | 212 | 513 | 88 | 194 | 37 | 223 | 366 | 338 |
| 35 | 210 | 536 | 93 | 204 | 39 | 238 | 381 | 367 |
| 38 | 218 | 555 | 97 | 212 | 40 | 248 | 394 | 390 |
| 41 | 225 | 563 | 98 | 218 | 40 | 257 | 404 | 409 |
| 43 | 232 | 581 | 99 | 225 | 40 | 263 | 414 | 424 |

| | | | | | | | | |
|---|---|---|---|---|---|---|---|---|
| 46 | 240 | 598 | 99 | 231 | 41 | 271 | 423 | 439 |
| 49 | 233 | 611 | 97 | 236 | 44 | 276 | 431 | 450 |
| 51 | 241 | 627 | 96 | 242 | 48 | 281 | 440 | 462 |
| 55 | 247 | 630 | 94 | 247 | 52 | 285 | 445 | 474 |
| 58 | 252 | 642 | 91 | 250 | 59 | 286 | 449 | 475 |
| 60 | 254 | 659 | 91 | 255 | 67 | 292 | 459 | 487 |
| 61 | 264 | 657 | 93 | 262 | 77 | 299 | 470 | 501 |
| 63 | 274 | 677 | 94 | 270 | 96 | 306 | 481 | 516 |
| 65 | 285 | 696 | 96 | 277 | 101 | 313 | 492 | 530 |
| 67 | 293 | 692 | 102 | 275 | 113 | 320 | 496 | 542 |
| 68 | 304 | 714 | 102 | 284 | 132 | 325 | 503 | 544 |
| 70 | 325 | 747 | 111 | 298 | 156 | 340 | 525 | 567 |
| 73 | 348 | 779 | 121 | 313 | 171 | 356 | 547 | 592 |
| 76 | 371 | 789 | 130 | 328 | 187 | 370 | 569 | 617 |
| 79 | 401 | 792 | 135 | 337 | 206 | 384 | 591 | 624 |
| 80 | 425 | 812 | 144 | 351 | 218 | 399 | 611 | 647 |
| 83 | 457 | 805 | 153 | 365 | 237 | 414 | 631 | 671 |
| 86 | 507 | 857 | 162 | 406 | 250 | 429 | 652 | 695 |
| 89 | 507 | 837 | 171 | 406 | 262 | 444 | 672 | 719 |
| 92 | 507 | 832 | 179 | 424 | 277 | 463 | 692 | 742 |
| 95 | 507 | 839 | 187 | 483 | 288 | 477 | 714 | 765 |
| 99 | 507 | 840 | 195 | 515 | 298 | 493 | 735 | 786 |
| 102 | 507 | 835 | 203 | 523 | 307 | 507 | 757 | 809 |
| 104 | 507 | 837 | 211 | 529 | 316 | 522 | 777 | 831 |
| 107 | 507 | 838 | 220 | 530 | 325 | 536 | 796 | 855 |
| 109 | 507 | 851 | 227 | 552 | 336 | 551 | 815 | 878 |
| 112 | 507 | 861 | 235 | 572 | 355 | 570 | 831 | 902 |
| 113 | 507 | 861 | 243 | 595 | 364 | 590 | 841 | 926 |
| 114 | 507 | 861 | 251 | 601 | 373 | 602 | 858 | 950 |
| 115 | 507 | 861 | 256 | 603 | 378 | 618 | 860 | 971 |

#### 2.2 Kinetics Fitting

A modified stacked sigmoidal function (Equation 1), based on [4] double sigmoid, was used for modeling $CH_4$ volumetric production in time,

$$V_{CH_4}^{STP}(t) = V_{CH_4}^{max} \cdot \left( \frac{p}{1+e^{\left(\frac{4r_1 \cdot (t_1 - t)}{V_{CH_4}^{max} \cdot p}\right)}} + \frac{1-p}{1+e^{\left(\frac{4r_2 \cdot (t_2 - t)}{V_{CH_4}^{max} \cdot (1-p)}\right)}} \right) \text{(Equation 1)}$$

Where $V_{CH_4}^{STP}$ is the specific $CH_4$ production in time (NmL$CH_4$ g $VS^{-1}$), $V_{CH_4}^{max}$ is the maximum specific volumetric production reached in the experiment (NmL$CH_4$ g $VS^{-1}$), $p$ is the proportion between ordinates values of 1st and 2nd stacked sigmoid, $t_1$ and $t_2$ are the time which production of the 1st and 2nd sigmoidal pattern reaches the maximum rate (d), $r_1$ and $r_2$ are the maximum rate of $CH_4$ production for the 1st and 2nd sigmoidal pattern, respectively (NmL$CH_4$ g$VS^{-1}$ $d^{-1}$),

Table 7, $CH_4$ production estimated by the kinetic model in Experiment 1

| Time (days) | Vinasse (NmL$CH_4$ g$VS^{-1}$) | Deacetylation Liquor (NmL$CH_4$ g$VS^{-1}$) | Filter Cake (NmL$CH_4$ g$VS^{-1}$) | Co-digestion (NmL$CH_4$ g$VS^{-1}$) | Celullose (NmL$CH_4$ g$VS^{-1}$) |
|---|---|---|---|---|---|
| 1 | 50 | 3 | 12 | 68 | 6 |
| 2 | 55 | 3 | 13 | 73 | 6 |
| 3 | 61 | 4 | 14 | 78 | 7 |
| 4 | 67 | 4 | 16 | 83 | 8 |
| 5 | 73 | 4 | 17 | 88 | 9 |
| 6 | 79 | 5 | 19 | 94 | 10 |
| 7 | 86 | 5 | 21 | 100 | 11 |
| 8 | 93 | 6 | 23 | 106 | 13 |
| 9 | 100 | 7 | 25 | 112 | 14 |
| 10 | 108 | 7 | 27 | 119 | 16 |
| 12 | 122 | 9 | 32 | 134 | 19 |
| 14 | 137 | 11 | 38 | 150 | 23 |
| 16 | 151 | 13 | 45 | 167 | 28 |
| 18 | 164 | 15 | 53 | 186 | 34 |

| | | | | | |
|---|---|---|---|---|---|
| 20 | 175 | 19 | 61 | 206 | 40 |
| 23 | 190 | 25 | 76 | 238 | 50 |
| 26 | 201 | 33 | 91 | 273 | 61 |
| 29 | 210 | 43 | 107 | 309 | 71 |
| 30 | 212 | 47 | 113 | 321 | 75 |
| 33 | 218 | 61 | 129 | 358 | 84 |
| 35 | 221 | 73 | 139 | 382 | 89 |
| 38 | 224 | 94 | 153 | 418 | 96 |
| 41 | 227 | 120 | 166 | 452 | 101 |
| 43 | 228 | 140 | 173 | 474 | 104 |
| 46 | 230 | 175 | 182 | 504 | 108 |
| 49 | 232 | 215 | 190 | 532 | 110 |
| 51 | 234 | 244 | 194 | 548 | 111 |
| 55 | 239 | 307 | 202 | 578 | 113 |
| 58 | 245 | 355 | 207 | 596 | 115 |
| 60 | 251 | 387 | 210 | 607 | 116 |
| 61 | 255 | 403 | 212 | 612 | 117 |
| 63 | 266 | 434 | 217 | 621 | 119 |
| 65 | 279 | 463 | 223 | 629 | 122 |
| 67 | 297 | 489 | 231 | 637 | 129 |
| 68 | 307 | 502 | 236 | 640 | 134 |
| 70 | 329 | 525 | 248 | 646 | 148 |
| 73 | 366 | 556 | 274 | 654 | 182 |
| 76 | 402 | 581 | 306 | 660 | 226 |
| 79 | 432 | 602 | 338 | 666 | 261 |
| 80 | 440 | 608 | 348 | 667 | 269 |

Table 8, CH4 production estimated by the kinetic model in Experiment 2

| Time (days) | Vinasse ($NmLCH_4$ $gVS^{-1}$) | Deacetylation Liquor ($NmLCH_4$ $gVS^{-1}$) | Filter Cake ($NmLCH_4$ $gVS^{-1}$) | Co-digestion ($NmLCH_4$ $gVS^{-1}$) | Cellulose ($NmLCH_4$ $gVS^{-1}$) | Filter Cake + vinasse ($NmLCH_4$ $gVS^{-1}$) | Filter Cake + Deacetylation Liquor ($NmLCH_4$ $gVS^{-1}$) | Vinasse + Deacetylation Liquor ($NmLCH_4$ $gVS^{-1}$) |
|---|---|---|---|---|---|---|---|---|
| 1 | 50 | 74 | 4 | 30 | 1 | 33 | 41 | 78 |
| 2 | 56 | 84 | 5 | 34 | 1 | 37 | 47 | 83 |
| 3 | 61 | 96 | 6 | 38 | 1 | 40 | 55 | 89 |
| 4 | 67 | 109 | 8 | 43 | 2 | 44 | 63 | 95 |
| 5 | 74 | 123 | 10 | 48 | 4 | 49 | 73 | 101 |
| 6 | 80 | 138 | 12 | 53 | 7 | 53 | 85 | 108 |
| 7 | 87 | 155 | 15 | 59 | 12 | 58 | 98 | 115 |
| 8 | 94 | 172 | 18 | 66 | 17 | 64 | 112 | 123 |
| 9 | 102 | 191 | 21 | 72 | 22 | 69 | 128 | 131 |
| 10 | 109 | 210 | 25 | 79 | 24 | 75 | 145 | 139 |
| 12 | 124 | 250 | 34 | 94 | 26 | 89 | 180 | 156 |
| 14 | 139 | 291 | 44 | 109 | 27 | 103 | 215 | 174 |
| 16 | 153 | 331 | 54 | 123 | 27 | 118 | 248 | 192 |
| 18 | 165 | 367 | 63 | 137 | 28 | 133 | 276 | 212 |
| 20 | 177 | 400 | 71 | 150 | 28 | 148 | 299 | 231 |
| 23 | 191 | 441 | 79 | 166 | 29 | 169 | 324 | 260 |
| 26 | 203 | 472 | 84 | 179 | 30 | 189 | 341 | 288 |
| 29 | 211 | 495 | 86 | 188 | 31 | 205 | 353 | 314 |
| 30 | 213 | 502 | 87 | 191 | 31 | 210 | 357 | 322 |
| 33 | 219 | 519 | 88 | 198 | 33 | 224 | 366 | 345 |
| 35 | 222 | 529 | 89 | 202 | 34 | 231 | 372 | 360 |
| 38 | 225 | 543 | 90 | 208 | 36 | 241 | 380 | 380 |
| 41 | 228 | 556 | 91 | 213 | 39 | 250 | 389 | 398 |
| 43 | 229 | 566 | 91 | 216 | 41 | 256 | 396 | 410 |

| | | | | | | | | |
|---|---|---|---|---|---|---|---|---|
| 46 | 231 | 581 | 92 | 222 | 45 | 264 | 406 | 427 |
| 49 | 234 | 597 | 93 | 227 | 51 | 271 | 417 | 443 |
| 51 | 236 | 609 | 94 | 232 | 55 | 276 | 425 | 453 |
| 55 | 241 | 636 | 97 | 241 | 65 | 287 | 442 | 475 |
| 58 | 248 | 657 | 99 | 250 | 75 | 296 | 457 | 491 |
| 60 | 255 | 671 | 101 | 256 | 83 | 302 | 467 | 502 |
| 61 | 259 | 679 | 102 | 259 | 87 | 306 | 473 | 508 |
| 63 | 268 | 693 | 104 | 266 | 96 | 312 | 484 | 519 |
| 65 | 280 | 708 | 106 | 274 | 106 | 319 | 496 | 531 |
| 67 | 295 | 722 | 109 | 282 | 117 | 327 | 508 | 544 |
| 68 | 303 | 729 | 111 | 287 | 122 | 331 | 514 | 550 |
| 70 | 322 | 743 | 114 | 296 | 135 | 339 | 527 | 563 |
| 73 | 354 | 762 | 120 | 311 | 154 | 352 | 548 | 583 |
| 76 | 389 | 779 | 126 | 328 | 175 | 365 | 569 | 605 |
| 79 | 421 | 793 | 134 | 347 | 197 | 380 | 591 | 627 |
| 80 | 431 | 798 | 137 | 353 | 204 | 385 | 599 | 635 |
| 83 | 456 | 810 | 146 | 373 | 226 | 401 | 622 | 658 |
| 86 | 475 | 820 | 156 | 395 | 246 | 418 | 645 | 683 |
| 89 | 488 | 829 | 166 | 417 | 265 | 435 | 669 | 708 |
| 92 | 497 | 836 | 177 | 440 | 283 | 454 | 692 | 734 |
| 95 | 503 | 841 | 188 | 463 | 298 | 472 | 716 | 761 |
| 99 | 508 | 847 | 202 | 494 | 315 | 497 | 746 | 797 |
| 102 | 510 | 851 | 212 | 516 | 325 | 516 | 769 | 824 |
| 104 | 511 | 852 | 219 | 530 | 331 | 529 | 783 | 842 |
| 107 | 512 | 855 | 227 | 551 | 339 | 548 | 804 | 869 |
| 109 | 512 | 856 | 233 | 564 | 343 | 560 | 817 | 887 |
| 112 | 512 | 857 | 240 | 582 | 348 | 579 | 836 | 914 |
| 113 | 512 | 858 | 242 | 588 | 350 | 585 | 842 | 923 |
| 114 | 513 | 858 | 244 | 594 | 351 | 591 | 848 | 931 |
| 115 | 513 | 859 | 246 | 599 | 353 | 596 | 854 | 940 |

**ACKNOWLEDGEMENTS**

This work was supported by FAPESP Project (2018/09893-1), FAPESP project (2016/16438-3), and FAPESP-BBSRC (2015/50612-8), The authors gratefully acknowledge the support of the Laboratory of Environment and Sanitation (LMAS) at the School of Agricultural Engineering (FEAGRI/UNICAMP), the National Laboratory of Biorenewables (LNBR/CNPEM), and the Interdisciplinary Center of Energy Planning (NIPE/UNICAMP),

#### 3. CONFLICTS OF INTERESTS

The authors have no relevant financial or non-financial interests to disclose,

#### 4. CREDIT AUTHOR STATEMENT

Maria Paula C, Volpi: Conceptualization, methodology, d ata curation

Brenno V, Lima: Methodology, data curation

Gustavo Mockaitis: Methodology, data curation mathematical modeling,

Bruna S, Moraes: Conceptualization, Review & Editing, Supervision,

#### 5. References

[1] J,M, Triolo, L, Pedersen, H, Qu, S,G, Sommer, Biochemical methane potential and anaerobic biodegradability of non-herbaceous and herbaceous phytomass in biogas production, Bioresour, Technol, 125 (2012) 226–232, https://doi,org/10,1016/j,biortech,2012,08,079,

[2] W, APHA, AWWA, Standard Methods for the Examination of Water and

Wastewater, twenty-sec, Washington, DC,, 2012,

[3] VDI 4630, Fermentation of organic materials, Characterization of the substrate, sampling, collection of material data, fermentation tests, Düsseldorf: Verein Deutscher Ingenieure, 2006,

[4] G, Mockaitis, G, Bruant, S,R, Guiot, G, Peixoto, E, Foresti, M, Zaiat, Acidic and thermal pre-treatments for anaerobic digestion inoculum to improve hydrogen and volatile fatty acid production using xylose as the substrate, Renew, Energy, 145 (2020) 1388–1398, https://doi,org/10,1016/j,renene,2019,06,134,
