## Supplementary Materials for "Use of lignocellulosic residue from second-generation ethanol production to enhance methane production through co-digestion"

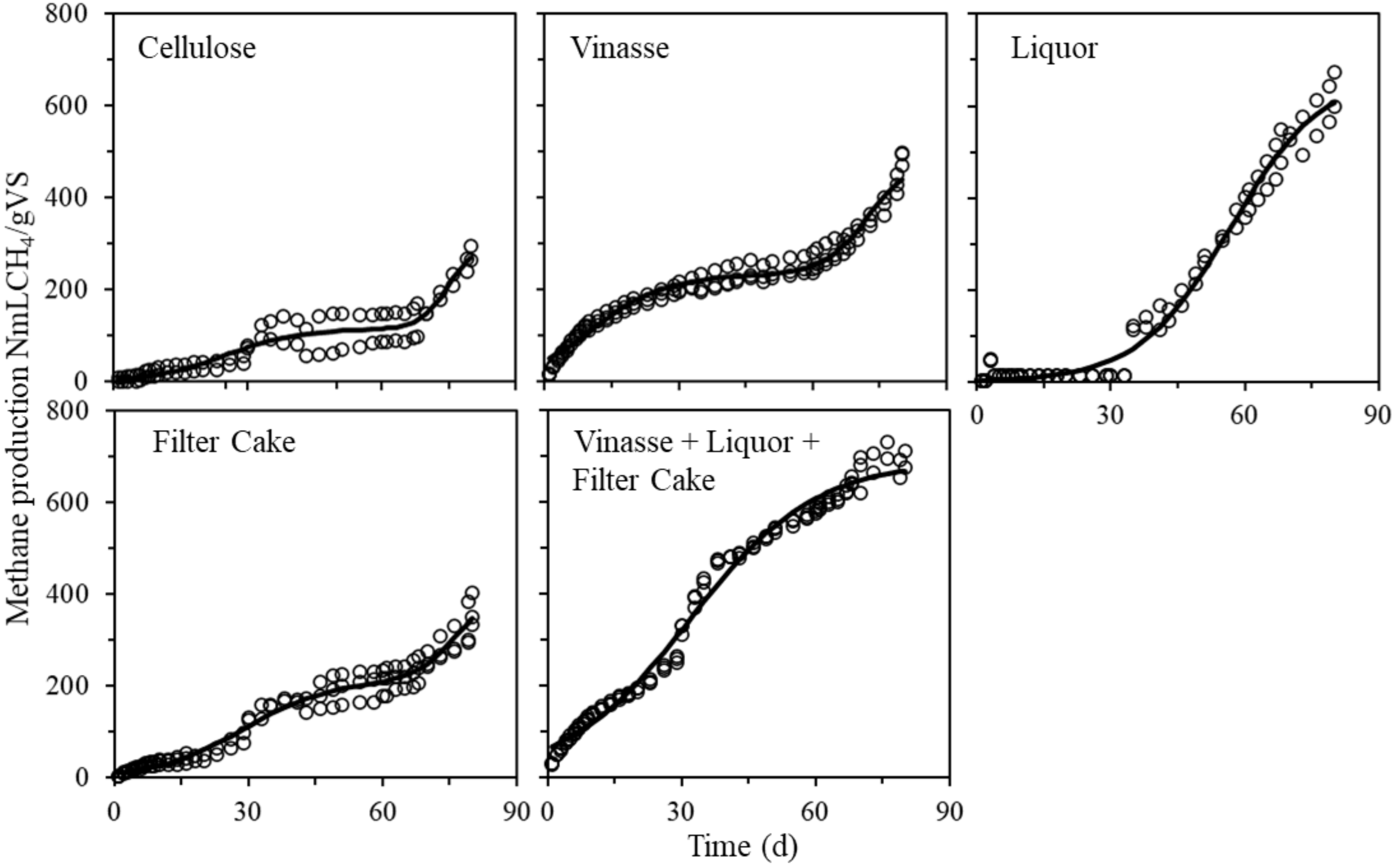


Figure 1SM – Time profiles for specific $CH_4$ production for each substrate and their combination for co-digestion, using sugarcane mill sludge as inoculum. Continuous line depicts the kinetic model fit to experimental data.

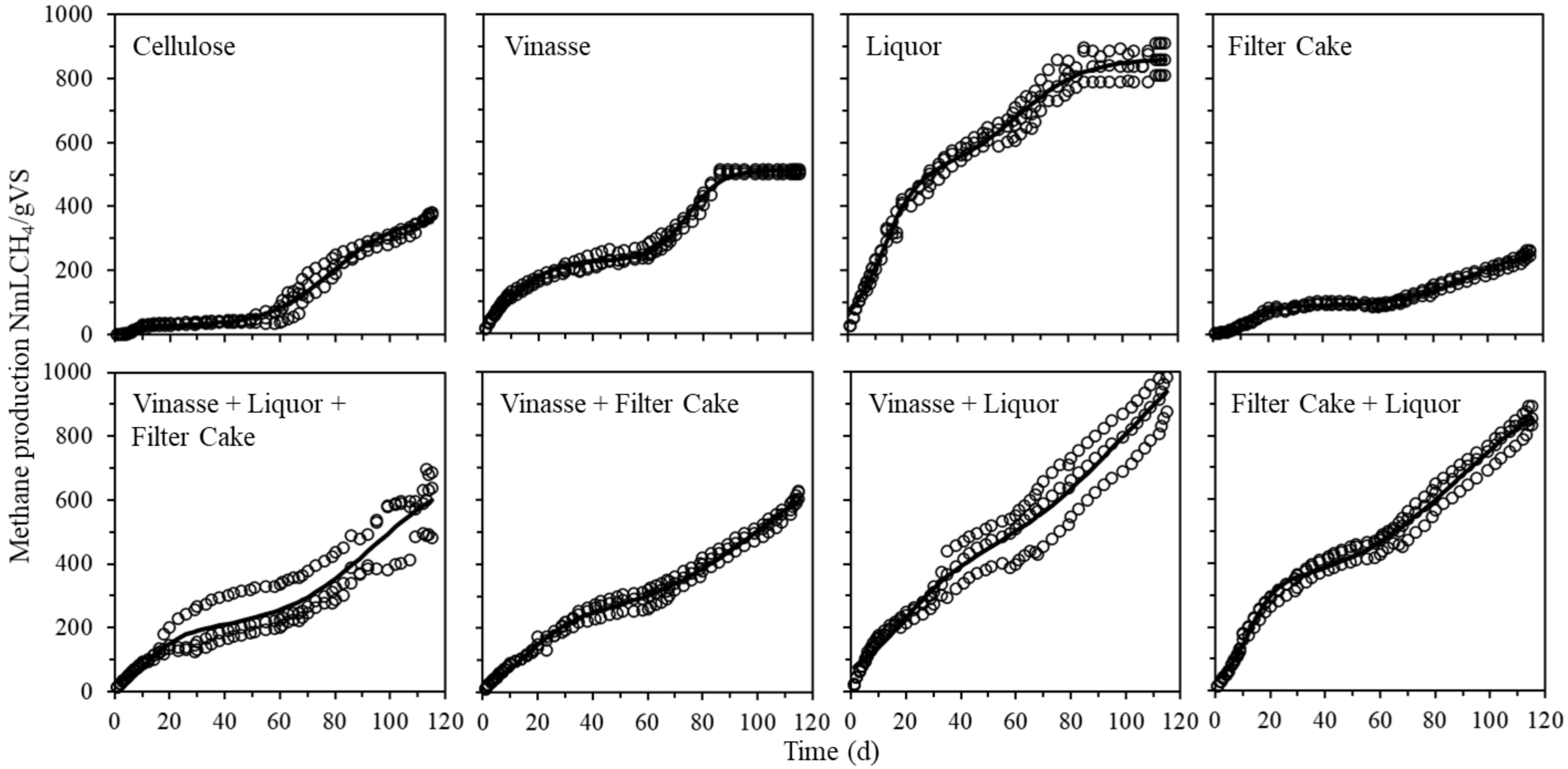


Figure 2SM – Time profiles for specific methane production for each substrate and their combination for co-digestion, using poultry slaughterhouse sludge as inoculum.
